# An arousal-gated visual circuit controls pursuit during *Drosophila* courtship

**DOI:** 10.1101/2020.08.31.275883

**Authors:** Tom Hindmarsh Sten, Rufei Li, Adriane Otopalik, Vanessa Ruta

## Abstract

Long-lasting internal states, like hunger, aggression, and sexual arousal, pattern ongoing behavior by defining how the sensory world is translated to specific actions that subserve the needs of an animal. Yet how enduring internal states shape sensory processing or behavior has remained unclear. In *Drosophila*, male flies will perform a lengthy and elaborate courtship ritual, triggered by activation of sexually-dimorphic P1 neurons, in which they faithfully follow and sing to a female. Here, by recording from males as they actively court a fictive ‘female’ in a virtual environment, we gain insight into how the salience of female visual cues is transformed by a male’s internal arousal state to give rise to persistent courtship pursuit. We reveal that the gain of LCt0a visual projection neurons is strongly increased during courtship, enhancing their sensitivity to moving targets. A simple network model based on the LCt0a circuit accurately predicts a male’s tracking of a female over hundreds of seconds, underscoring that LCt0a visual signals, once released by P1-mediated arousal, become coupled to motor pathways to deterministically control his visual pursuit. Furthermore, we find that P1 neuron activity correlates with fluctuations in the intensity of a male’s pursuit, and that their acute activation is sufficient to boost the gain of the LCt0 pathways. Together, these results reveal how alterations in a male’s internal arousal state can dynamically modulate the propagation of visual signals through a high-fidelity visuomotor circuit to guide his moment-to-moment performance of courtship.

---

Across the animal kingdom, complex and lengthy courtship rituals are interposed between mate recognition and copulation, during which behavior towards conspecifics is altered. In *Drosophila*, courtship typically commences after the male performs a discrete chemosensory assessment in which he extends a foreleg to sample a female’s pheromones and evaluate her suitability as a sexual partner^1,2^. The initiation of courtship represents a discrete switch in a male’s behavior, in which he transitions from being apathetic or “blind” to a female to vigorously tracking her while singing a species-specific song. This enduring pursuit is thought to be essential to offer choosy females the time to assess between alternative mates, and to entice her to copulate^3^. Given the temporal and metabolic investment of lengthy courtship rituals, however, aroused males must still remain sensitive to behavioral and sensory feedback to prevent continued pursuit of an inappropriate or unreceptive mate. A male’s courtship pursuit must therefore be both persistent and flexible to maximize chances of reproductive success.

*Drosophila* courtship is largely mediated by ∼1500 neurons marked by expression of the sex-determination factor *fruitless* (*fru*^*M*^)^4–6^, offering exceptional traction to understanding its underlying circuit logic. Recent work has identified a population of −20 sexually-dimorphic P1 neurons as a central node within the Fru^M^+ courtship circuitry that regulates a male’s entry into courtship^7–10^. P1 neurons receive convergent input from excitatory and inhibitory chemosensory pathways that tune their activity to the pheromones of a conspecific female mate^11–14^. Moreover, transient activation of P1 neurons drives sustained courtship displays, including visual pursuit and singing, even towards inanimate objects^11,13,15–17^. P1 neurons thus appear to gate the release of an enduring arousal state in which the salience of female sensory signals is transformed. How does this long-lasting internal state restructure sensorimotor circuits to transform a female from an indifferent visual object to a target of desire?

**Courtship reflects a dynamic arousal state**

To explore how neural activity is altered by a male’s arousal state, we developed a virtual reality visual system based on previous, integrated hardware design (J.L. Weisman and G. Maimon, personal communication and ref. 18; **Fig. 1a, Extended Data Fig. 1**) in which a tethered male can court a simplified fictive ‘female’ represented as a high-contrast dot projected onto a conical screen. Activation of P1 neurons expressing the light-gated cation channel CsChrimson induced males to chase the autonomously moving visual target over many meters in this 2D virtual world, while performing the ipsilateral wing-extensions typical of courtship song production (**Fig. 1b, Extended Data Fig. 2, Supplementary Video t**). Males maintained the target at close range within the center of their visual field, replicating the oriented pursuit characteristic of free courtship behavior (**Fig. 1c-d, Extended Data Fig. 2, Supplementary Video 2**). Wild-type males did not display any overt behavioral responses (**Fig. 1c, Extended Data Fig. 2**), underscoring that courtship of this minimal ‘female’ is driven by P1 neuron activation.

**Figure 1.**
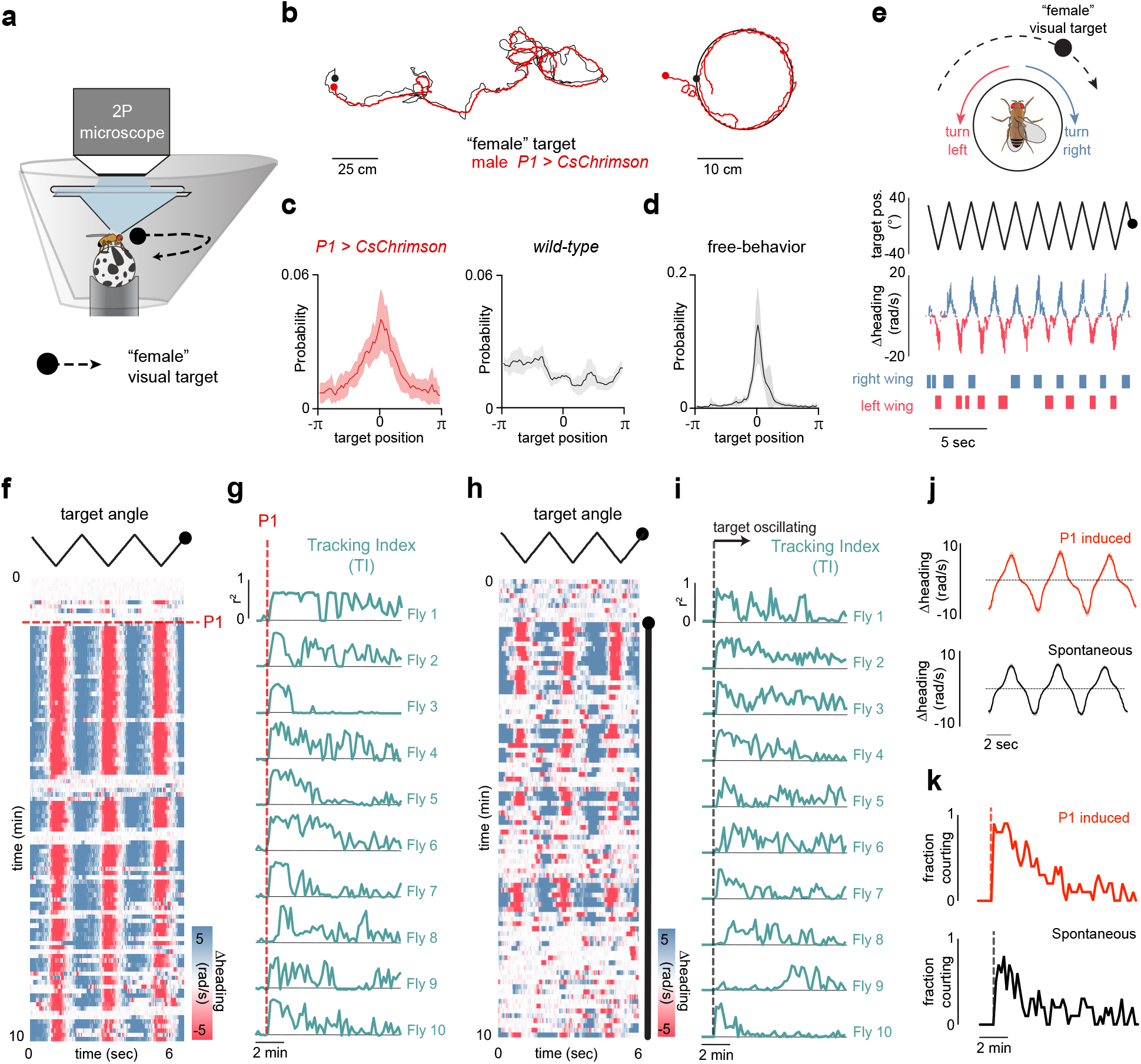
P1 neurons release a dynamic state of sexual arousal. **a**, Schematic of behavioral setup used for tethered courtship. **b**, 2-D paths of males in closed-loop courting a randomly moving target (left) or a target moving in circle (right) over the course of 200 seconds during activation of P1 neurons. **c**, Distribution of angular positions of the target female relative to males expressing CsChrimson in P1 neurons or wild-type animals during tethered courtship. **d**, same as (**c**) but for freely courting pairs of wild-type animals. **e**, Example of a courting male’s tracking during courtship relative to the position of the target stimulus. Raster at bottom indicates times of unilateral extensions of the male’s wings. **f**, Example of a male’s turning behavior during a courtship trial, before and following 3-second optogenetic activation of P1 neurons expressing CsChrimson (red line). Each row consists of three stimulus cycles, with target angle of the visual stimulus shown at top. **g**, Bivariate correlation between the stimulus position and male turning (Tracking Index) for l0 flies following 3-second optogenetic activation of P1. Top trace is the same animal as in (**f**). **h**, Same as **f** but for a spontaneously initiated courtship trial. Black bar indicates when the visual stimulus is present and oscillating in front of the male. **i**, same as **g**, but for trials where courtship was spontaneously initiated. **j**, Average turning response during courtship in trials where courtship was induced by activation of P1 neurons versus trials where courtship was spontaneously initiated. **k**, Fraction of males actively engaged in courtship over the course of a trial in optogenetic trials and spontaneously initiated trials. Dashed lines indicate LED onset (red) or the onset of visual motion (black). All shaded line plots are mean±s.e.m. Details of statistical analyses and sample sizes are given in Supplementary Table 1.

The dynamics of a male’s arousal has been challenging to elucidate in the context of natural social encounters, where continuously changing sensory feedback from another fly could contribute to its regulation. We reasoned that monitoring changes in a male’s response to an invariant visual stimulus should offer an overt behavioral readout of his internal state, dissociated from changes in his external sensory environment. We therefore presented males with a dot which traversed back and forth along an arc with a constant angular velocity, and whose size subtended on the male retina approximated that of a female during natural courtship. Tethered males were initially indifferent to this visual stimulus but after a brief (3 sec) optogenetic activation of P1 neurons began to faithfully track and sing to it for many minutes (**Fig. 1e-g, Supplementary Video 3**). To quantify how the fidelity of a male’s courtship unfolded over time, we defined a “tracking index”-the time-averaged squared correlation between the angular position of the visual stimulus and a male’s turning speed. Examining the tracking indices of different males revealed that, while all courted persistently after transient P1 activation, the structure of their courtship was highly idiosyncratic (**Fig. 1g)** as males did not incessantly court the visual target following P1 neuron activation. Rather, their pursuit fluctuated in both fidelity and vigor over a trial and was frequently punctuated by brief pauses during which they ceased tracking despite the continued presence of the visual stimulus (**Fig. 1f-g**).

Males thus transition between two distinct states following transient P1 neuron activation: an aroused state in which the visual profile of the ‘female’ target elicits turning, and a latent state in which it does not. Males in this latent state, however, remain primed to reinitiate pursuit, in contrast to unaroused males prior to P1 activation. Over time, the probability that a male occupies this aroused state steadily decreases, suggesting that the priming effect of transient P1 activation subsides (**Fig. 1k**). Males can maintain this latent drive to court even in the absence of continuous sensory input or behavior as they rapidly resumed pursuit if the visual target was briefly (30 sec) removed and then reintroduced (**Extended Data Fig. 3**).

Persistent courtship of the virtual target could also be triggered by allowing males to taste the pheromones on a conspecific female, replicating his sensory assessment of a prospective mate^1,13,14^ (**Extended Data Fig. 3**). However, while pheromone pathways converge onto P1 neurons to initiate courtship, these chemical cues are not essential to arouse a male^19,20^. Indeed, females lacking cuticular pheromones remain highly attractive targets, underscoring that chemical communication in *Drosophila* serves largely to hone a male’s inherent visual arousal, promoting courtship towards conspecific females and suppressing tracking of inappropriate mates^19–21^. We therefore asked whether persistent courtship pursuit could be evoked by visual input alone. As males failed to track the oscillating target in the absence of P1 neuron activation, we modified the visual stimulus to more closely mimic the natural statistics of female motion such that the stimulus continued to rotate at a constant angular velocity, but now also appeared to recede and advance from the tethered male by changing in angular size, replicating the inter-fly distance of a courting pair (**Extended Data Fig. 4, Supplementary Video 4**). We also socially isolated males to enhance their courtship drive^11,16,19,22^. Males spontaneously initiated courtship towards this translating ‘female’ stimulus (**Fig. 1h-i**), displaying the same persistent and faithful pursuit, unilateral wing-extensions, and idiosyncratic bout structure characteristic of courtship evoked by exogenous activation of P1 neurons (**Figure 1i-k, Extended Data Fig. 4, Supplementary Video 5**). We observed no behavioral responses in tethered females (n=9, data not shown), consistent with sexually dimorphic nature of courtship pursuit. Visual cues are therefore sufficient to release a persistent state of arousal in *Drosophila* males, providing an inroad to examine how courtship-related arousal is represented within P1 neurons.

## Pt neurons reflect arousal during courtship

We next performed functional calcium imaging of P1 neurons as males initiated spontaneous courtship pursuit, allowing us to relate their activity to a male’s changing arousal (**Fig. 2a**). We observed that P1 neurons became robustly activated at the onset of courtship, even when males initiated pursuit many seconds after presentation of the visual stimulus (**Fig. 2b-d**, time to initiation: 0.3-314 sec). P1 activity remained elevated throughout the duration of a courtship bout, returning to baseline each time that males temporarily paused their pursuit (**Fig. 2b-c**,**f**). As a consequence, fluorescence changes in P1 neurons were tightly correlated with a male’s tracking index (**Fig. 2b-e**, r = 0.562±0.092, mean±st.d.). In contrast, P1 activity displayed a weak relationship to a male’s linear (r = 0.231±0.099, mean±st.d. **Fig. 2h**) or angular velocity (r = 0.296±0.071, mean±st.d., **Fig. 2i**), likely because males must run and turn to track the target, and was uncorrelated with the visual stimulus (r = 0.004±0.006, mean±st.d., **Fig 2g**). The activity of P1 neurons thus more closely aligns with a male’s courtship state than the sensorimotor implementation of his pursuit.

**Figure 2.**
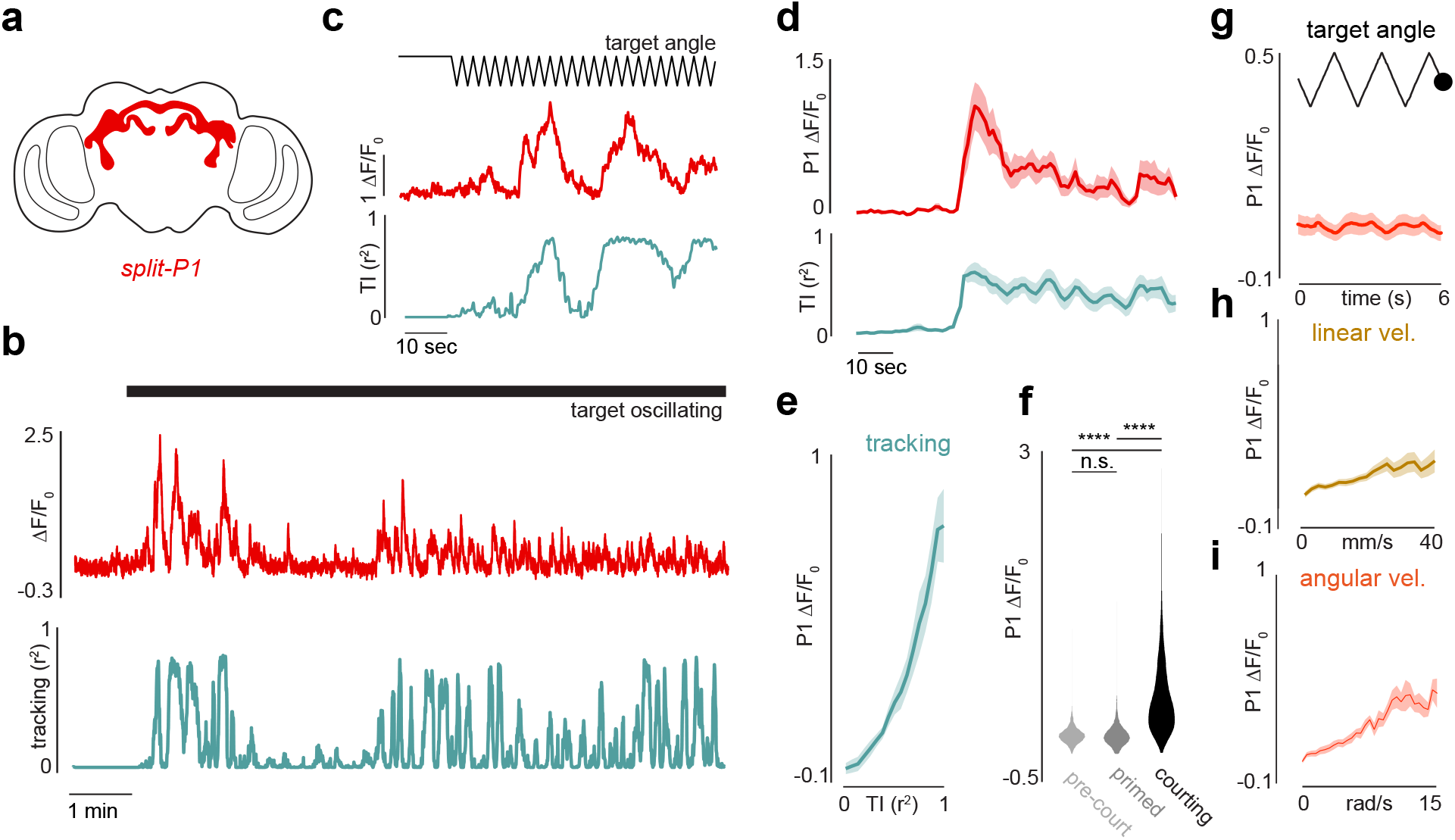
P1 neurons reflect courtship arousal. **a**, Schematic of the anatomy of P1 neurons in the male brain. **b**, Example of P1 activity during a spontaneous courtship trial. Top trace: functional response (LF/F0) of P1 neurons expressing GCaMP over the course of a courtship trial; bottom trace: tracking index over the course of the trial. Black line indicates when the visual stimulus is oscillating. **c**, Zoomed-in view of the onset of courtship in (**a**). **d**, Top: Average response of P1 neurons aligned to the onset of courtship; bottom: average tracking index aligned to the onset of courtship. **e**, Functional responses (average LF/F0) of P1 neurons versus behavioral tracking index. **f**, Distributions of P1 activity (LF/F0) before courtship was initiated, during periods where the animals did not track the visual target but remained primed to reinitiate courtship, and during active courtship. **g**, Functional responses (average LF/F0) of P1 neurons aligned to the cycles of the visual target. **h**, Functional responses (average LF/F0) of P1 neurons versus the male’s linear velocity. **i**, same as **h**, but for the male’s angular velocity. All shaded line plots are mean±s.e.m.; n.s. indicates p > 0.05; **** indicates p <0.000l. Details of statistical analyses and sample sizes are given in Supplementary Table 1.

Recording from Fru+ neurites within the lateral protocerebral complex (LPC), a neuropil richly innervated by the P1 neurons^7,10^, revealed activity that was similarly correlated with the intermittent bout structure of courting males (**Extended Data Fig. 5**). To confirm that activity in these higher-order Fru+ neurons, including the P1 population, correlates with a male’s arousal state but not his motor actions, we used wide-field motion to evoke optomotor responses and mimic the alternating turns of courtship pursuit in males that were sexually unaroused. No relationship was observed between changes in LPC fluorescence and the angular or linear velocity of unaroused males executing optomotor reflexes (**Extended Data Fig. 5**).

Notably, calcium signals in both P1 neurons and the LPC decayed between courtship bouts, when males ceased tracking but remained primed to reinitiate pursuit, suggesting that this latent state must be stored subcellularly or in the activity patterns of other downstream neurons (**Fig. 2f, Extended Data Fig. 5**). P1 neurons therefore reflect the distinct timescales over which courtship is regulated, with activity time-locked to the entry into a lasting internal state and ongoing fluctuations that correspond to bouts of arousal during which the male tracks the target. To transform these representations of a male’s arousal state into behavior, however, they must impinge on sensorimotor circuitry to acutely regulate a male’s pursuit. Indeed, males initiate tracking of a female only upon becoming sexually aroused^1,15-17^, indicating that visual pathways must be functionally coupled to the motor circuits controlling steering behavior in a highly state-dependent manner.

## Courtship arousal modulates the gain of LCt0a visual projection neurons

LCl0a visual projection neurons were recently identified as small spot motion detectors essential for accurate tracking during courtship^23^, making them an intriguing neural locus to investigate arousal-dependent processing of female cues. The LC10a population is comprised of ∼100 neurons per hemisphere, the majority of which are Fru+, that convey retinotopically organized visual signals from the lobula to the medial anterior optic tubercle (AOTu)^23,24^. To confirm that LCl0a neurons are necessary to track a fictive female target in tethered courtship^23^, we selectively expressed the light-gated anion channel GtACR1 in a subset of this population and used the spatial precision of a two-photon microscope to inhibit their axon terminals within the AOTu of a single hemisphere (**Fig. 3a**). Unilateral silencing of LC10a terminals selectively reduced turning towards the visual stimulus when it traversed through the ipsilateral but not the contralateral hemifield (**Fig. 3b-c**). LC10a neurons are thus required for faithful pursuit in both tethered and free males^23^.

**Figure 3.**
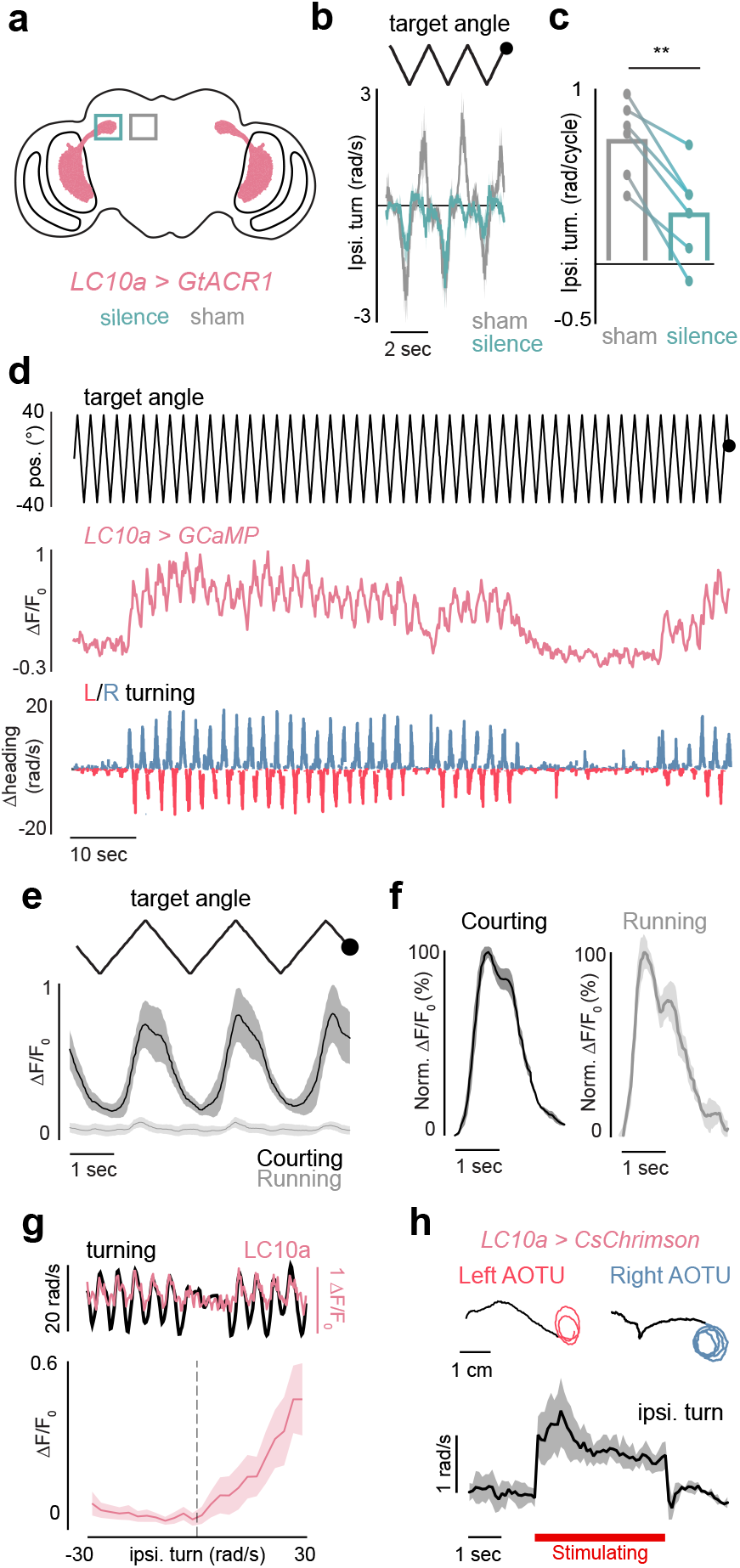
Modulation of LCt0a neurons during courtship. **a**, Schematic of LCl0a neurons expressing GtACRl and ROIs used for silencing (or sham-silencing) in a single hemisphere. **b**, Example of the average turning of a male to the visual stimulus during silencing of LC10a neurons in the right hemisphere versus sham trials. **c**, Average ipsilateral turning during LC10a silencing versus sham trials. **d**, Representative example of LCl0a neuron responses in the right AOTu during courtship bouts. From top to bottom: angular position of the target stimulus; functional responses (LF/F0) of LCl0a neurons expressing GCaMP; angular velocity of male. **e**, Functional responses (average LF/F0) of LC10a neurons expressing GCaMP during periods of courtship versus periods of running without tracking, aligned to the visual stimulus position. **f**, Average peak-normalized responses (LF/F0) of LC10a neurons during courtship versus during running. **g**, Top: example of LC10a responses versus animal turning. Bottom: Average activity (LF/F0) of LC10a neurons versus the magnitude of ipsiversive turning. **h**, Top: sample 2D path of male before (black) and during optogenetic stimulation of LC10a neurons in the left (red) or right (blue) hemisphere. Bottom: Average evoked ipsilateral turning during unilateral stimulation of LC10a neurons expressing CsChrimson. All shaded line plots are mean±s.e.m.; n.s. indicates p > 0.05; ** indicates p < 0.01; *** indicates p <0.001. Details of statistical analyses and sample sizes are given in Supplementary Table l.

To examine whether LC10a neuron visual responses were altered by a male’s arousal, we monitored GCaMP activity in their axon terminals within the AOTu as a male engaged in spontaneous courtship pursuit. LC10a neurons only weakly responded to the visual target at baseline, prior to courtship initiation. Once courtship pursuit commenced, however, LC10a axon terminals were robustly activated each time the visual target swept across the male’s ipsilateral hemifield (**Fig. 3d-e**). The shape of LC10a responses remained largely unchanged, pointing to alterations in the gain and not the tuning properties of this visual pathway (**Fig. 3f**). The strength of LC10a signaling therefore appeared strongly dependent on a male’s internal state, such that this neural population reliably encoded the motion of the target only when males were highly aroused (**Fig. 3d-e**). Notably, during the brief epochs where courting males temporarily stopped tracking LC10a neurons returned to their low baseline activity level (**Fig. 3d**), underscoring that the gain of these pathways is modulated on a moment-to-moment timescale during courtship pursuit. The gain enhancement of LCl0a neurons was specific to epochs when males were actively tracking and not simply running or turning in response to widefield motion (**Fig. 3e, Extended Data Fig. 6**) revealing that this heightened sensitivity was distinct from more the generalized changes in arousal associated with locomotion^25,26^. To further dissociate the gain of LC10a neurons from the motor implementation of visual tracking, we took advantage of the fact that males cannot turn left and right simultaneously. We introduced a second target to males that were actively courting, whose position was equal and opposite to the first target, yielding identical stimulation simultaneously to both eyes (**Extended Data Fig. 6**). LC10a neurons continued to strongly respond in these aroused males, even when the male failed to turn, or turned in the contralateral direction (**Extended Data Fig. 6**). These experiments highlight that the elevated gain of LC10a neruons is independent of the motor execution of a male’s pursuit.

The tight correspondence between the amplitude of LC10a visual responses and the magnitude of a male’s ipsilateral turns (**Fig. 3g**) supports that these neurons do not simply represent visual objects during tracking, but actually underlie a male’s faithful pursuit. Consistent with previous work^23^, we found that unilateral activation of LC10a axon terminals in the AOTu was sufficient to drive robust ipsilateral turning for the duration of stimulation (**Fig. 3h**). Anterograde trans-synaptic tracing using *trans-*Tango^27^ revealed that the downstream synaptic partners of LC10a neurons have axon terminals in the lateral accessory lobe (LAL) of the fly brain (**Extended Data Fig. 7**), a region densely innervated by descending neurons (DNs)^28^. Connectomic analysis in females^29^ supports that LC10a neurons connect to relatively few downstream neuropils, with LAL innervating neurons representing a major target (91.2% of identified LCl0a neurons are pre-synaptic to LAL neurons, **Extended Data Fig. 7**). Together, these results suggest that sexual arousal gates the propagation of sensory signals through a concise sensorimotor circuit.

## A network model predicts visual pursuit during courtship

To examine whether modulating the gain of LC10a neurons could capture the dynamics of visual pursuit during courtship, we constructed a simple network model mimicking the architecture of this visuomotor circuit. We modeled the LCl0a neurons as a population of integrate-and-fire units, each of which covered a portion of the male’s visual field (estimated as 270**°** with 15**°** of binocular overlap^29^) and utilized the motion-based receptive fields estimated by Ribeiro *et al*. to structure excitatory input to these model neurons (**Fig. 4a**)^23^. The total number of spikes of the LC10a population in the right and left hemisphere was integrated and compared by downstream units such that the strength of ipsilateral turning depended on the difference in firing rate between hemispheres in a given time-bin, approximating lateralized input to descending pathways. Importantly, the model was sensitive to the change in the angular position of moving objects, but not their angular size on the retina, consistent with the broad tuning of LC10a receptive fields and behavioral evidence that males tracked targets of varying size with equivalent vigor (**Extended Data Fig. 4**)^23^.

**Figure 4.**
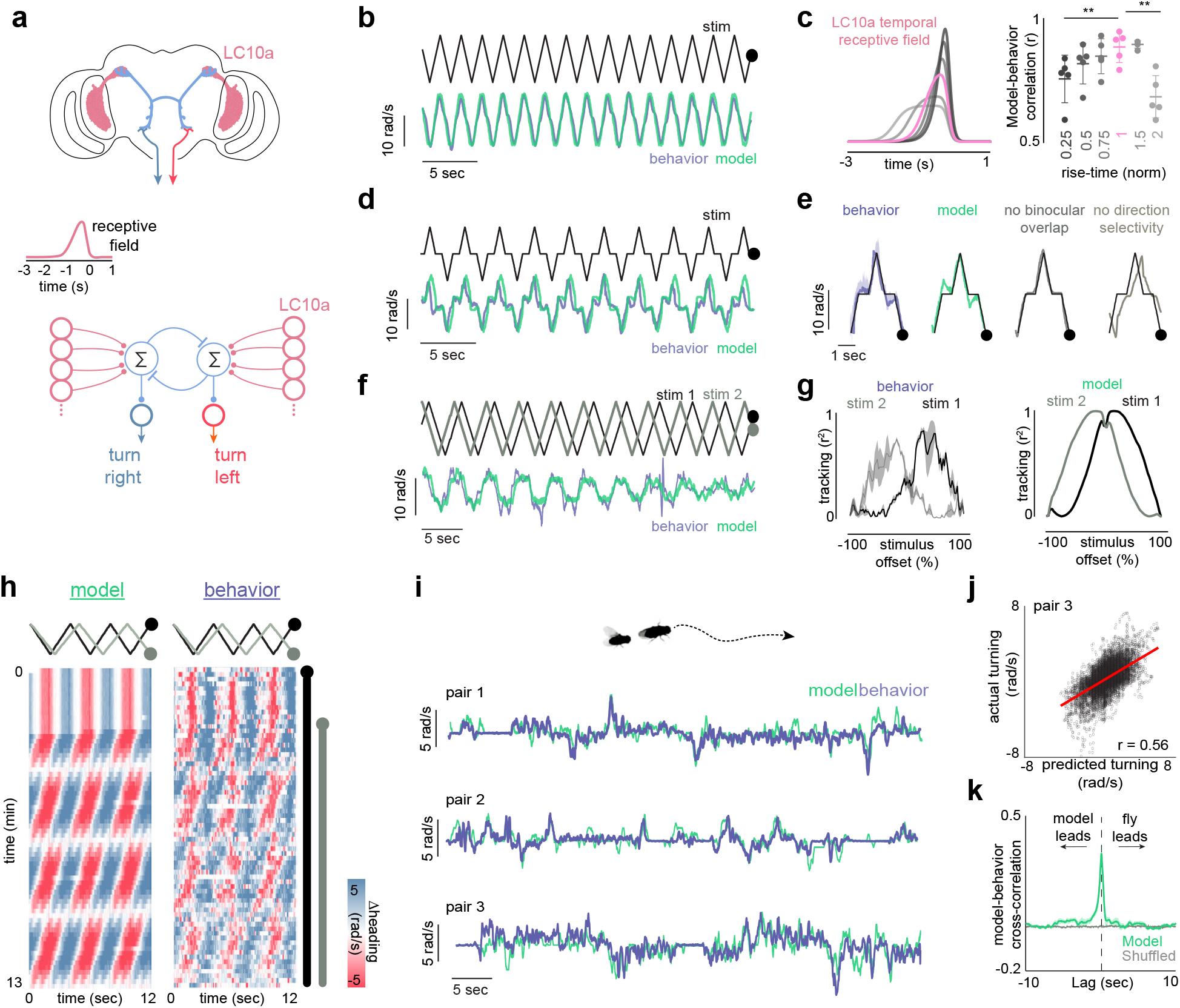
A network model of LCt0a neurons recapitulates visual pursuit during courtship. **a**, Top: Schematic of putative circuit anatomy from LCl0a to descending neurons; middle: LCl0a temporal receptive fields; bottom: schematic of network model. **b**, Predicted versus actual turning response to an invariant target during activation of P1 neurons. **c**, Left: normalized LCl0a receptive fields with varying rise-times (K, see methods). Right: correlation between predicted and actual responses to the simple stimulus in **b** for the receptive fields shown to the left. **d**, Example of predicted versus actual turning response to a single target that pauses for 500ms on each stimulus cycle **e**, From left: average turning response to a single stimulus cycle; predicted response from full model to a single stimulus cycle; predicted response of a model with no binocular overlap; predicted response of a model not selective for progressive versus regressive motion. **f**, Example of predicted versus actual turning of two targets with a drifting phase-offset. **g**, Left: Average tracking index of the first target (black) and second target (grey) versus the phase-offset between the targets. Positive values indicate that the first stimulus leads in phase. Right: same as left but based on the model’s predictions. **h**, Example of predicted versus actual turning response to two targets with a drifting phase-offset (as in **f**,**g**) across the courtship trial. Black line indicates when first target is present, grey line indicates when the second target is present. **i**, Examples of predicted versus actual turning responses of males in freely courting pairs of flies over the course of l minute. **j**, Predicted versus actual male turning over the course of a courtship trial for all frames, red line denotes the linear fit (r = 0.56). **k**, average cross correlation between predicted and actual turning during free courtship behavior. All shaded line plots are mean±s.e.m.; **indicates p < 0.01. Details of statistical analyses and sample sizes are given in Supplementary Table l.

We initially investigated which parameters of the model were necessary to accurately predict the sensorimotor transformation of an aroused male in response to a single fictive female target (**Fig. 4b**). We found that small deviations in the rise-time and shape of the model’s spatiotemporal receptive fields strongly degraded its performance, indicating that the specific temporal tuning properties of LC10a neurons are essential for the model in order to replicate the faithful pursuit of a courting male (**Fig. 4c**). Thus, despite the fact that LCl0a receptive fields were measured from the attenuated responses of unaroused males, they optimally predicted courtship tracking, further supporting that state-dependent modifications to visual processing reflect alterations in the gain of LCl0a signaling without necessitating alterations to the parameters of visual detection.

While previous recordings suggested that LC10a neurons were relatively insensitive to the direction of motion for a visual stimulus^23^, we found that selectivity to progressive (front-to-back) motion was necessary for the model to accurately track the visual target in a time-locked fashion (**Fig. 4b, Extended Data Fig. 8**). To behaviorally test this prediction, we slowed the visual target to temporally resolve progressive and regressive motion, and found that males turned ipsilaterally to the stimulus target location during progressive motion, whereas their turning slowed during periods of regressive motion (**Extended Data Fig. 8**). Likewise, imaging from Fru+ neurons within the AOTu confirmed that LCl0a neurons were direction selective in aroused animals (**Extended Data Fig. 8**). Thus, consistent with the model’s prediction, both neural and behavioral responses display directional tuning during courtship, a property that should enhance the accuracy of a male’s courtship pursuit by triggering him to turn to a female only as she moves away from the center of his field-of-view and not towards it.

Behavioral evidence from houseflies suggests that males may be able to predict the motion of a target as it passes from one visual field to another during courtship, enhancing the fidelity and continuity of pursuit^31^. Given the direction-selectivity we observe, we reasoned that this anticipatory behavior may arise from the small region of space where the receptive fields of LCl0a neurons from both hemispheres overlap, allowing a moving target to evoke turning before it has crossed a male’s central axis. To assess the contribution of binocular overlap to courtship pursuit, we modified the oscillating visual target so that it briefly (500ms) paused in front of the male each time it traversed across his field of view. Our model predicts that as this stimulus enters the region of binocular overlap, it will preferentially excite contralateral LC10a neurons tuned to progressive motion, driving males to turn in the contralateral direction as the target stops, before it has crossed his central axis. Consistent with this prediction, aroused males robustly turned in the contralateral direction as the stimulus paused, closely replicating the magnitude and timescale of the model’s overshoot (**Fig. 4d**, r = 0.86, p<0.00001, Pearson correlation between model and behavior; **Fig. 4e**). Removing either the binocular overlap of LC10a neurons or their direction selectivity from the model degraded its performance, highlighting how both are critical to capture the turning dynamics of courting males.

A central element of the model is that the difference in activity between LC10a neurons in the right and left hemispheres dictates the direction and strength of a male’s turns. To explore this feature, we introduced a second visual target that oscillated at 98% of the velocity of the first target, allowing us to systematically measure a male’s steering behavior as the phase relationship between the targets continuously drifted over time (**Supplementary Video 6**). Our model predicts that when the two targets are closely aligned in phase, males will preferentially turn in response to the leading target, due to the directional tuning of LC10a neurons and interhemispheric competition, generating relatively selective pursuit of only one target (**Fig. 4f-h**). However, as the two targets drift further apart, a male’s turning response should become less well-aligned to either stimulus and once the two targets were completely out-of-phase, high fidelity tracking should cease altogether as the two eyes receive identical input (**Fig. 4f-h, Extended Data Fig. 6**). Aroused males presented with this complex stimulus responded nearly indistinguishably from the model, exhibiting similar tracking on a moment-to-moment basis as the offset of the two targets shifted, albeit real males were slightly more selective in their pursuit of just one fictive female target (**Fig. 4g**,**h;** r = 0.83, p<0.00001, Pearson correlation between model and behavior).

Given the model’s predictive power, we asked whether it could replicate the dynamics of freely moving males engaged in natural courtship, where female trajectories are inherently more circuitous and variable. Remarkably, using just the relative positional information of the female, the model accurately predicted the steering behavior of courting males over many minutes (**Fig. 4i-k, Extended Data Fig. 9** mean r = 0.52±0.06, n = 6 pairs, p < 0.00001 for all flies). Furthermore, by assuming that model males match their linear velocity to that of the female, we could simulate the high-fidelity tracking of a freely-courting male within a circular arena (**Supplemental Video 7**). The properties of this simple visuomotor circuit are thus largely sufficient to account for the faithful tracking of a female, supporting the essential role of this visual pathway for courtship pursuit (**Fig. 3**).

## P1 neurons dynamically regulate pursuit during courtship

Our model supports that LCl0a visual signals become tightly coupled to descending pathways that control steering behavior during courtship, allowing them to predictably drive turning in response to a moving target. Yet, males only engage in courtship pursuit after becoming aroused, indicating that this visual pathway lies dormant until P1 neurons are activated. Moreover, even following the initiation of courtship, males alternate between bouts of vigorous pursuit and brief pauses during which the visual profile of the ‘female’ target fails to elicit a turning response. Although P1 neurons do not directly innervate the AOTu, the tight correspondence between their activity and ongoing changes to a male’s arousal (**Fig. 2**) suggests that signals about the intensity of this internal state may be propagated to LC10a neurons to modulate their gain and give rise to the intermittent bouts of courtship pursuit. In accord with this possibility, we found that optogenetic activation of P1 neurons acutely enhanced the gain of LCl0a signaling, driving LC10a neurons to robustly respond each time the visual target swept across a male’s field of view, yielding incessant and almost invariant tracking (**Fig. 5a-e**). P1 activity is thus sufficient to regulate the propagation of visual information through the central brain by controlling the gain of the LC10a visual signaling.

**Figure 5.**
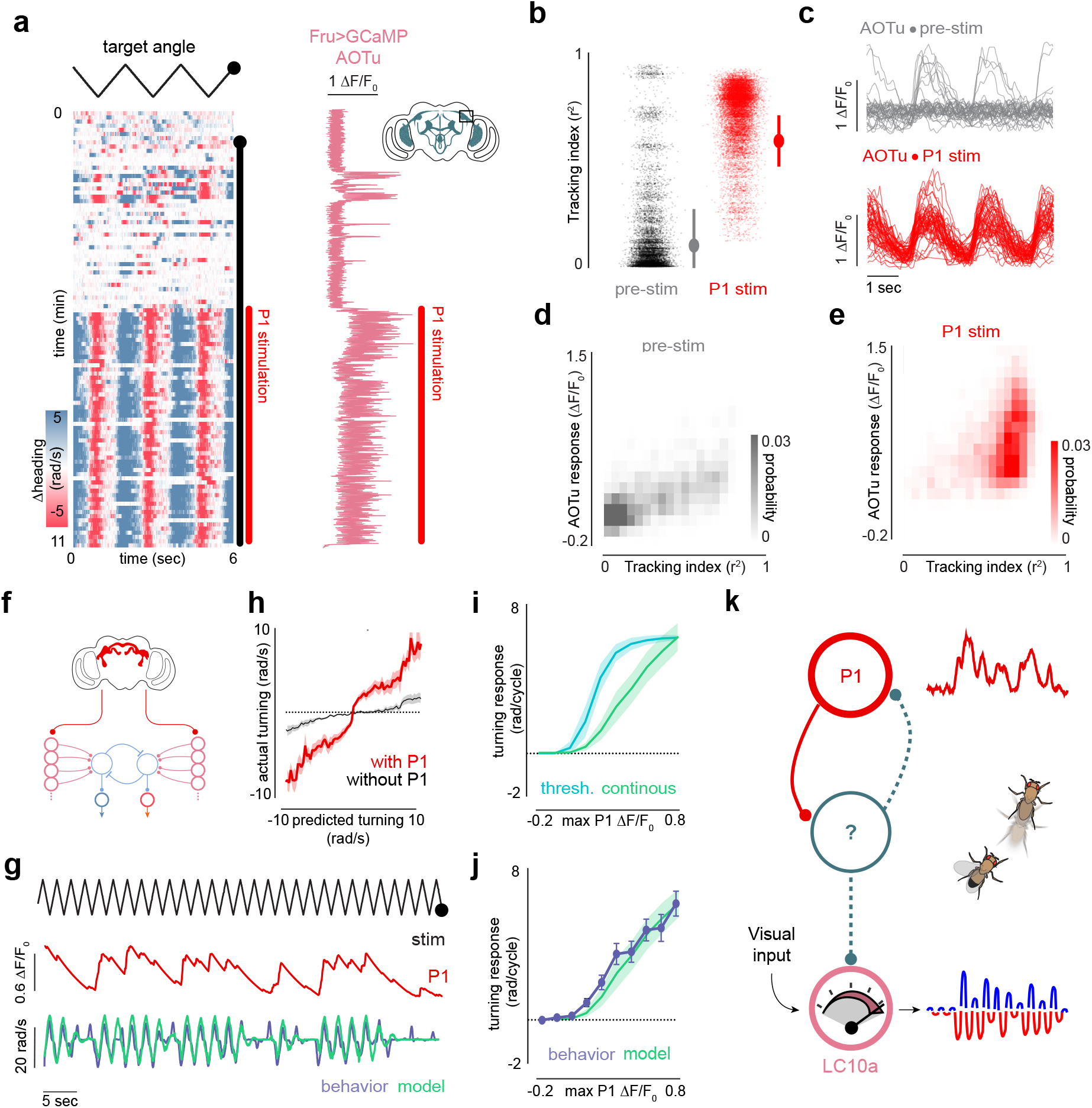
P1 neurons acutely reflect and regulate arousal and pursuit. **a**, Left: Example of a male’s turning over the course of 4 minutes of spontaneous courtship followed by 6 minutes of optogenetic activation of P1 neurons expressing CsChrimson (red line). Black line indicates when the visual stimulus is oscillating. Right: functional response (LF/F) of putative LC10a Fru^+^ neurons in the AOTu expressing GCaMP during the trial. **b**, Distribution of the tracking index during spontaneous courtship versus during P1 activation for the animal in (**a**). **c**, Functional responses (LF/F0) of putative LCl0a neurons in the AOTu for each three-cycle period during spontaneous courtship (top) and during P1 activation (bottom) for the animal in (**a-b**). **d**, Density plot of tracking index versus functional responses (LF/F0) of putative LCl0a neurons in the AOTu on each stimulus cycle during spontaneous courtship across animals. **e**, same a (**g**) but during continuous activation of P1 neurons. **f**, Schematic of P1 model: input current to LCl0a is scaled by P1 activity (LF/F0). **g**, Example of i actual versus predicted turning using model incorporating the activity of P1 neurons. **h**, Actual versus predicted turning of courting animals for models with (red) and without (grey) incorporating P1 activity. **i**, Net ipsiversive turning responses during one stimulus cycle of a continuous model versus a threshold-based model as a function of P1 LF/F0. **j**, Same as **i** but with behavioral data shown in purple. **k**, Schematic of model for P1-mediated arousal dynamically regulating LCl0a gain during courtship. Details of statistical analyses and sample sizes are given in Supplementary Table 1.

Finally, we reasoned that if P1 neurons dynamically modulate the gain of signaling through the LC10a visuomotor circuit, incorporating P1 activity into our model should enhance its predictive power by accounting for fluctuations in a male’s arousal during spontaneous courtship. Strikingly, we found that simply scaling the net input current of LCl0a neurons by the normalized activity of P1 neurons (ΔF/F, **Fig. 5f**) significantly improved the accuracy of the model, yielding a faithful prediction of a male’s moment-to-moment turning (**Fig. 5g-h, Extended Data Fig. 10**). By including P1 activity, the model no longer incessantly responded to the visual stimulus, and instead accurately predicted the intermittent bout structure exhibited by courting males. Incorporating P1 activity increased the fraction of predicted turns that were accompanied by a real turn (9l.9± .6 of turns versus 6 .l±l .3 without P1 activity), while the fraction of real turns accompanied by a predicted turn remained high (93.7±6.08% of turns versus l00 without P1 activity **Extended Data Fig. 10**).

P1-mediated arousal could serve as a binary gate on the flow of visual information through the LCl0a sensorimotor circuit or it could instead continuously tune the strength of visual signaling in a graded manner. To differentiate between these possibilities, we modeled how the strength of ipsiversive turns depended on P1 neuron activity using a threshold-based versus a continuous-gain function (**Fig. 5i**). We found that the magnitude of turning responses in courting males exhibited a linear relationship to P1 neuron activity with a slope that closely matched a continuous model (**Fig. 5j**), in contrast to the sigmoidal relationship predicted by the threshold model. These results further support that a male’s sexual arousal is a scalable function that exists along a continuum of strengths, allowing for P1 neurons not only to trigger the discrete entry into a courtship state, but to also continuously regulate the strength of signaling through visuomotor circuits to shape ongoing behavior **(Fig. 5k)**.

## Discussion

Internal arousal states motivate distinct patterns of behavior over short and long timescales, allowing for the temporary emergence of innate behavioral programs like fighting, feeding, or mating, that subserve physiological needs^32–35^. In this study, we reveal how a male’s enduring state of sexual arousal transforms the salience of visual cues by engaging a latent visual circuit that guides the moment-to-moment implementation of courtship pursuit. Leveraging a male’s willingness to persistently track a simple fictive ‘female’ stimulus allowed us to dissociate changes in male’s sensory environment from changes in his internal state and gain insight into how a male’s arousal rapidly patterns his ongoing courtship pursuit. We find that even in response to an unchanging visual stimulus, males display striking variation in their moment-to-moment tracking behavior, pointing to either intrinsic variation in the fidelity of their visuomotor circuits or fluctuations in their drive to chase the visual target. Our work supports the latter, with the idiosyncratic pursuit of males mediated by a highly reliable visuomotor circuit whose activity is continuously regulated by P1 neuron-mediated arousal, which functions like a rheostat to control the propagation of visual signals in a graded manner. Indeed, exogenous activation of P1 neurons boosts the gain of LCl0a neurons, driving males to become unwavering in the fidelity of their courtship pursuit, such that their behavioral responses to a visual target are almost fully specified by the properties of LC10a circuit.

The sensorimotor implementation of visual tracking thus appears to be largely independent from the drive to perform it, paralleling the concept of how a fixed and reflexive motor pattern can be unleashed via an innate releasing mechanism from classic ethology^32,35^. This segregated circuit logic reconciles two competing needs for mate pursuit and discrimination: first, to allow for the robust and faithful tracking necessary to entice a conspecific female to copulate and, second, to maintain sensitivity to sensory and behavioral feedback, allowing a male to flexibly control his pursuit on a moment-to-moment timescale. Given the lack of visual discrimination apparent in aroused males^18^, continuous feedback is required to hone a male’s pursuit towards only receptive conspecific females. For example, males will promiscuously chase a variety of moving targets yet transiently terminate their pursuit if they detect pheromones carried by heterospecific females or other males^12,19,20,36^, underscoring how chemical cues play a key role in mate discrimination, by both gating the initial entry into courtship and tightly regulating its continued perpetuation. P1 neurons, as the site of convergence for both excitatory and inhibitory signals emanating from other flies^11–14,37,39^, offer a powerful nexus for such dynamic control. As inhibitory pheromone pathways directly impinge onto the P1 neurons^13,14^, they could acutely reverse the gain of LC10 pathways, rendering males temporarily “blind” to an inappropriate mate and suppressing futile pursuit, enabling them to reorient to more suitable sexual partners. Consistent with this possibility, acute silencing of P1 neurons by either direct optogenetic inhibition or via activation of ascending inhibitory pheromone pathways temporarily interrupts male courtship^38,39^. Moreover, males deficient in their ability to detect cuticular pheromones court heterospecific females vigorously but indiscriminately^12,20,36,40^, pointing to the critical importance of chemical cues to temper a male’s inherent attraction to moving targets. Likewise, sampling excitatory pheromones on a conspecific female during courtship would reactivate P1 neurons and reinvigorate a male’s pursuit, preventing the decay in arousal we observe in tethered males offered only a visual target.

In the wild, *Drosophila* meet and mate on fermenting fruits where diverse species will frequently congregate^41^. Our work highlights multiple properties of the LCl0a visuomotor circuit that would facilitate the identification and faithful pursuit of a conspecific female within this complex social environment: (i) LC10a neurons are tuned to small moving targets^23^ over a range of angular sizes (**Extended Data Fig. 4**), enabling males to track females from a variety of distances; (ii) the directional tuning and binocular overlap of LC10a visual pathways allows males to smoothly track a female and anticipate her future position even before she has crossed the center of his field-of-view (**Fig. 4d-e**); (iii) interhemispheric competition, likely implemented through contralateral inhibition, enables males to maintain selective pursuit of a single female, suppressing turning to other flies crossing into their visual field (**Fig. 4f-h**); (iv) the LC10a circuit propagates visual signals in a highly state-dependent manner, preventing males from aberrantly responding to visual objects except in an appropriate behavioral context (**Fig. 3**); (v) finally, the persistence of P1-evoked arousal state provides an enduring representation of the presence of a suitable sexual partner, motivating males to continue to sample and pursue target flies to mate. How this persistent state is maintained remains to be determined in particular as P1 calcium levels are not sustained between courtship bouts, when males remain primed to reinitiate their pursuit, suggesting this latent drive to court is encoded elsewhere in the circuit. For example, the persistent activity of recently identified pCD neurons^38^ may comprise part of a Fru+ network that could perpetuate a male’s arousal state and potentially feedback to the P1 population to encode ongoing fluctuations in arousal. Alternatively, this latent drive could be stored by subcellular molecular cascades in P1 neurons themselves, regulating their intrinsic excitability following transient activation.

Behaviorally relevant visual features in *Drosophila* are thought to be encoded by multiple distinct and spatially segregated populations of visual projection neurons^23,24^, including LC10a, which form optic glomeruli within the central brain. Our work suggests how a male’s arousal state may orchestrate the different motor components of a male’s courtship display by taking advantage of this modular system. Following P1 neuron activation, we observed that a male’s pursuit of the invariant ‘female’ stimulus consistently outlasted his willingness to sing to it (data not shown), suggesting that these two behaviors can be elicited by the same visual cue but display distinct sensitivity to the male’s slowly waning arousal. Moreover, while acute silencing of LC10a neurons inhibits accurate tracking, singing remains intact^23^, indicating that different populations of visual projection neurons guide the different behavioral components of courtship, each with distinct visual tuning and sensitivity to a male’s arousal. The anatomically segregated nature of optic glomeruli renders them a logical substrate for this modular control^42^, allowing for state-dependent coupling of visual features to different motor actions. Such a modular organization could underlie the flexible patterning of behavior, allowing the same internal state to differentially regulate distinct visually-guided behaviors, or the same visual pathways to be differentially used in distinct social contexts, like pursuit of potential mates versus competitors during courtship and aggression^42–44^. This modularity could also facilitate the evolution of male courtship displays, allowing for the rapid diversification of the structure and sequence of a male’s courtship ritual^2^ by tinkering with either the tuning of a visual pathway or its dependence on a male’s arousal state. The dynamic gating of sensorimotor circuits we describe offers a general mechanism for state-dependent behavior, honing the expression of different behavioral programs throughout the life of an individual or across different species.

## DATA AVAILABILITY

All data underlying this study available upon request from the corresponding author.

## CODE AVAILABILITY

The code underlying the generation and analysis of the network model is available upon request from the corresponding author.

## ACKNOWLEDGEMENTS

We thank J. Biden for providing hope in the fight against fascism. We thank K. Kuchibhotla, B. Noro, L. Abbott, and R. Coleman for comments on the manuscript; I. Ribeiro for sharing the LC10a split-GAL4 driver line, and Y. Aso for LexAop-GCaMP6s/UAS-CsChrimson flies; J. Weisman and G. Maimon for generously sharing their designs for an integrated, projector-based system for presenting tethered, walking flies with visual stimuli in advance of publication; R. Coleman, J. Petrillo, I. Morantte, and A. Siliciano for technical advice; and G. Maimon, C. Bargmann, L. Abbott, and members of the Ruta laboratory for discussion. This work was supported by the Simons Collaboration to the Global Brain, a Reem-Kayden Award, and 5R35NS111611 from the National Institute of Health to V.R., a David Rockefeller Fellowship to T.H.S, and an HHMI Hanna H. Gray Fellowship to A.O. This material is based upon work supported by the National Science Foundation Graduate Research Fellowship Program under Grant No. l9 6 29. Any opinions, findings, and conclusions or recommendations expressed in this material are those of the author(s) and do not necessarily reflect the views of the National Science Foundation.

## AUTHOR CONTRIBUTIONS

T.H.S. and V.R. conceived of and designed the study. T.H.S. and R.L. performed tethered behavioral experiments. A.O. carried out the free courtship assays in Fig. 4.T.H.S. performed and analyzed functional imaging experiments. T.H.S. designed and implemented the model. T.H.S. and V.R. analyzed data and wrote the manuscript with input from R.L and A.O.

## METHODS

### Data reporting

Preliminary experiments were used to assess variance and to optimize behavioral conditions. Experiments were not randomized, and the experimenters were not blind to conditions. For tethered courtship assays, unless otherwise noted, only experiments during which animals exhibited courtship towards the visual targets were included for analysis, selected based on a tracking index > 0.5 for at least 1 second and the presence of at least one unilateral wing extension. For modelling of free behavior, only flies that courted for >90% of the time before copulation were included.

### Fly stocks and husbandry

Flies were housed under standard conditions at 25°C on a l2h light-dark cycle, except for flies expressing channelrhodopsin variants which were dark-reared. Fly stocks used were as follows: *Split-P1 GAL4* (D. Anderson, California Institute of Technology), *LC10-SS1 GAL4* (I. Ribeiro and B. Dickson, Janelia Research Campus), *UAS-GtACR1* (A. Claridge-Chang, Duke-NUS), *UAS-CsChrimson, LexAop-GCaMP6s* (Y. Aso, Janelia Research Campus), *20X-UAS-CsChrimson*.*mVenus* (V. Jayaraman, Janelia Research Campus), *Fru-LexA* (B. Dickson, Janelia Research Campus), *R;Trans-Tango* (G. Barnea, Brown University). The following stocks were obtained from the Bloomington Drosophila Stock Center: CantonS, *20XUAS-IVS-jGCaMP7f* (BDSC 79031), *20XUAS-IVS-GCaMP6s* (BDSC 42746), *OL0019B* (BDSC 68336).

Supplementary table 1 provides detailed descriptions for all genotypes used in each experiment.

### Free behavior assays

All assays were performed with male and virgin female flies 3-5 days post-eclosion. Flies were isolated -l2 hours post-eclosion and reared with flies of same sex at low density (5-10 animals) in food vials (d = 3cm, h = 9cm) for 3-4 days. Courtship assay chambers were custom-milled bowls with a 40mm diameters, 3mm depths, and sloped edges to prevent flies from walking upside-down. Chambers were covered with a thin sheet of clear acrylic. Flies were added to the chamber by aspiration without anesthetization and videos were then recorded for 30 minutes with a Point Grey FLIR Grasshopper USB3 camera (GS3-U3-32S4M-C: 3.2 MP, l2l FPS, Sony IMX252, Monochrome) using the Fly-capture2 Software Development Kit (FLIR). Videos were captured from underneath the chamber at 40 frames per second and with a resolution of 34 ± 1 pixel per micron. All behavioral assays were conducted in a heated, humidified room (25 °C, 46% average relative humidity) on a backlit surface (Logan Electric Slim Edge Light Pad A-5A, 5 00K, 6 klx). The x-y coordinates and orientations of the male and the female flies were tracked using the Caltech FlyTracker^45^. The angular position of the female target in the male’s visual field was calculated with a custom MAT-LAB (Mathworks) script wherein, for each frame, the male and female x-y coordinates and orientations were translated and rotated such that the male was situated at the origin, facing zero degrees. The angular position of the female was then calculated as the inverse tangent of her x coordinate over her y coordinate in this new basis.

### Fly tethering and dissection

Flies used for closed-loop behavioral assays were briefly (<30 sec) anaesthetized on CO_2_, and subsequently tethered to a stainless-steel insect pin (size 00, *d=*0.3mm, *l=*4cm, Fine Science Tools) by their thorax. Anaesthetized flies were placed on a custom-milled plate and held in place by a short string across the neck, and an insect pin was brought in and centered on the back of the thorax. A small dollop of UV-curable clue (Loctite AA 3106) was manually placed at the contact point and cured with a UV gun for 0.5 sec. Flies were left to recover from anesthesia for 1-4 hours in a dark chamber humidified by a small wet paper towel.

For open-loop behavioral assays and two-photon functional imaging, flies were briefly anaesthetized on CO_2_ and tethered to a custom-milled plate similar to those used in previous studies^12,46^. Flies were held in place by a string across the neck and fixed to the holder by both eyes and the back of the thorax using UV-curable glue (Loctite 3106). To minimize brain motion during functional imaging, the proboscis was also glued to the mouthparts. The string was subsequently removed, and flies were left to recover in a warm, humidified chamber (25°C, 50-70% humidity) in the dark. For behavioral experiments, flies were transferred to the ball after 2-6 hours. For functional imaging experiments, flies were left in the dark until immediately before the assay, at which point the cuticle was removed to give optical access to the central brain without anesthesia. The tethering plate was filled with saline (l08mM NaCl, 5mM KCl, 2mM CaCl_2_, 8.2 mM MgCl_2_, mM NaHCO_3_, 1 mM NaH_2_PO_4_, 5 mM trehalose, 10 mM sucrose, 5 mM HEPES, pH .5 with osmolarity adjusted to 265 mOsm) to cover the fly’s head, and the cuticle between the eyes was cut with a 30-gauge needle and removed with forceps. The trachea covering the top of the central brain was removed from both hemispheres with forceps. Flies were subsequently transferred to the ball and left to recover in darkness for at least 30 minutes.

### Virtual reality preparation

For our virtual courtship preparation, we adapted an existing hard-ware design for presenting tethered flies with visual stimuli (Jazz L. Weisman and Gaby Maimon, *in preparation*). Male flies rested on a small 6.35mm diameter ball shaped from Last-A-Foam FR-6l8 (General P1astics)^47,48^ painted manually with uneven black spots using a Sharpie. The Styrofoam ball was held by a custom-milled aluminum base with a concave hemisphere of 6.75mm. A 1mm tract drilled through the base connected to air supplied at −0.8 L/min. The aluminum base was held in place by a custom printed (Carbon 3D) contraption. The ball was illuminated by infrared LED flood lights, and imaged with a Point Grey FLIR Firefly camera (FMVU-03MTM-CS) with a 9 mm/lx WD Video Lens (InfiniStix) by way of a mirror (ThorLabs #ME05-G01). The ball was surrounded by a 270° conical screen with a large diameter of ∼220 mm, a small diameter of ∼40mm, and a height of ∼60mm. The screen was cut from matte cardboard using a laser cutter, and fitted into a custom 3-D printed screen-holder with a tilted slit for placing and forming the screen shape (Extended Data Fig. 1).

The visual stimulus was projected around the male from a DLP 30l0 Light Control Evaluation Module (Texas Instruments). During optogenetic behavioral assays (Fig. 1b-h), light was projected by way of a 40×40cm mirror (First Surface Mirrors, custom cut) from above the fly, whereas it was rear-projected onto the front of the screen during all other behavioral experiments and for two-photon calcium imaging due to the sterics of the objective. The red and green LEDs in the projector were turned off using the DLP Display and Light Control EVM GUI (Texas Instruments), leaving only the blue LED. The lens of the projector was also covered by blue filter-paper. Together, this minimized any aberrant activation of neurons in animals expressing the light-gated cation channel CsChrimson, as *split-P1*>*CsChrimson* animals did not initiate courtship spontaneously when only the blue LED was on. As males walked on the spherical treadmill, all three rotational axes of the ball were read-out by the FicTrac2.0 software^49^ at 60Hz in real time. FicTrac was linked by a socket to MAT-LAB, which read out the estimated angular position of the ball at each frame. The projector received input from the same computer running FicTrac via an HDMI cable, and was controlled via a mini-USB cable to the same computer. Visual stimuli were generated in the MATLAB-based ViRMEn software^50^ and projected onto the screen using custom perspective transformation functions. On each frame of an experiment, the animal’s updated position was read-out in MATLAB, and either stored or used to transform the position of the visual stimulus in the case of closed-loop assays. The net visual refresh rate of the visual stimulus ranged from .6Hz to 58.9Hz. All other experimental variables were regulated in MATLAB on a single computer interfacing with an Arduino Uno controlling the LED. Wing-extensions were detectable via the same camera used for ball tracking, but a second camera (Point Grey, FMVU-03MTM-CS) was used for better visual access to the wings during manual scoring of wing-extensions.

### Closed loop behavioral assays

All flies used for closed-loop experiments were reared in the dark on sugar-yeast food to prevent low levels of ion-flux through light-gated ion channels during development, as previously described^12^. Male flies were transferred l-2 days after eclosion to food containing 400µM all-trans-retinal 48 h before behavioral assays^16^ and kept at low density (3-males/vial). Pin-tethered male flies were placed at the center of the ball and left to acclimate to the ball for 30-60 minutes in darkness before the experiment was started. This also served to bring male arousal to a baseline before optogenetic activation of P1 neurons. The change in the animal’s heading and the integrated x-y position of the ball was read out from FicTrac on each frame. This positional information was used to update the animal’s position and heading in the virtual ViRMEn world. Thus, when the fly turned clockwise, the world was rotated counter-clockwise, and vice versa, simulating the natural visuo-motor coupling of a freely behaving fly. Optogenetic stimulation was delivered by way of the red-LED in the projector (4µW/ mm^2^ at 600nm), which consistently drove animals to court. The visual stimulus in closed loop experiments either consisted of (i) a black dot that followed a stereotyped motion pattern (Fig. 1b, right) or (ii) a black dot that followed a random trajectory influenced by the male’s own motion (Fig. 1b, left). In both cases, the stimulus was presented on a white background with a light-grey floor (Supplemental Movie 2).

(i) For the former case, the diameter of the dot was such that it occupied ∼30° of the screen when positioned 10mm in front of the animal in the virtual world. This is larger than the size of a natural female at the same distance but allowed us to keep the female some distance away from the male in the virtual world while she could still exhibit a similar angular size to a close-up female. This extra distance between the male and female was important to prevent males from accidentally colliding with the female in virtual world, as there is no tactile feedback available. Distances in the virtual world were determined such that 1 radian of the ball the fly walked on was equivalent of one virtual unit. The object rotated in a circle around the virtual world, whose diameter was equivalent of 20cm, with a rotational velocity of -l0°/s.

(ii) For the latter case, the diameter of the dot was the same as during (i), but the female moved in an unpredictable and pseudorandom pattern. The female target originally moved in a random direction with a constant linear velocity of −30mm/s. At each point, she had 20 probability of switching her current heading direction to a new heading direction, drawn pseudorandomly from a normal distribution with its mean equaling the current heading and a variance of 35°. After a switch, she was prohibited from switching direction over the course of the next second. To prevent the male from losing the female in the infinite world, the female was softly-bounded around 120mm away from the male in all directions (her turns were biased towards the male’s current location). To prevent the female from being on top of the male, she was also softly-bounded 10mm from the male (her turns were biased away from the male’s current location). Experiments ranged from 10-30 minutes in duration, but, for consistency, only the first l0 minutes from each animal was included for further analysis.

### Open loop behavioral assays

#### Optogenetically induced courtship

All flies used for closed-loop experiments that expressed channelrhodopsin variants were reared in the dark on sugar-yeast food. Male flies were transferred l-2 days after eclosion to food containing 400µM all-trans-retinal 48h before behavioral assays and kept at low density (3-males/vial). After plate tethering, male flies were transferred to a humidified warm chamber to recover from anesthesia for 2-6h. Flies were subsequently placed on the center of the ball and left to acclimate for at least 1h. This ensured that any remnant arousal caused by activation of P1 neurons during mounting had ample time to decay, and that flies were in a baseline state at the onset of the trials. Two 1.5-mm optic fibers (Edmund Optics) were coupled to two high-power red LEDs (660nm, LED Engin) mounted on a heat-sink (Ohmite), and placed directly above the head of the fly. Optical activation of P1 neurons was made with a single optical pulse, yielding a net power 8µW/mm^2^ at 600nm. Since the power of the red and green LEDs in the projector was set to zero, flies were presented with a small dark blue dot (**ø** ∼28°, mimicking the angular size of a female fly 2 mm away from the male) on a light blue background. This dot oscillated in a symmetric 160° arc about the male, at a constant distance (i.e. size) and with a constant angular velocity.

For most single-stimulus experiments (see exception below), the visual target oscillated with a velocity of -l60°/s. Flies were presented with the visual stimulus for 60-l20s before optogenetic activation of P1 neurons. This allowed us to examine the flies’ baseline responses to the visual target, and ascertain that P1 neurons were not being activated by light from the projector as animals did not initiate tracking during this baseline period. P1 neurons were subsequently transiently activated by a single 3-s continuous optical pulse. Following initial P1 activation, we continued to monitor the animal’s motion in response to the visual stimulus for the remainder of the trial (between 9 to 29 minutes). For consistency, only the first l0 minutes from each animal was included for analysis.

For stop-and-go motion, the target ceased to move for 500ms at the center of the screen on each cycle, and subsequently continued on along its arc path. For estimating behavioral responses to progressive versus regressive motion, the angular velocity of the target was slowed down by 50% to 80°/s to give ample time for the expression both components of the behavior. For experiments where two dots of different velocity were presented, trials began with a 1-3 minute “blank” period where no stimulus was presented, after which a single stimulus appeared for 1-2 minutes. All males robustly courted the target. A second stimulus was subsequently added to the screen. This stimulus was identical to the first stimulus, but moved at 98 of the velocity (Figs. f-h). The velocity of the faster dot was 80°/s.

When examining model parameters, we continuously activated P1 neurons optogenetically during tethered behavioral experiments, allowing us to measure visual responses from males in a uniformly heightened arousal state (Fig. 4).

#### Pheromone induced courtship

Virgin male flies were collected following eclosion and group-reared at low density for 2-3 days before behavioral assays. To prepare the stimulating female abdomen, we removed the wings and legs from a 3-7-day old CantonS virgin female to ensure that the distal portion of the male’s forelegs could readily contact her abdomen. We subsequently manually attached a pin to her dorsal thorax, and attached this pin to a long custom 3-D printed holder designed to fit around the conical screen. The female abdomen was positioned ∼1.5cm and 90° to the left of the male before the trial was started, and viewed from the side using an IR-sensitive camera (Point Grey, FMVU-03MTM-CS) equipped with a 94-mm focal length lens (InfiniStix). As during optogenetic activation of P1 neurons, male flies were presented with the visual stimulus for several minutes before he was presented with the stimulating abdomen of a conspecific female, allowing us to internally control his baseline response to the visual target. The experimenter subsequently gently centered the abdomen in front of the male and brought it into contact with the male’s foreleg using a micromanipulator (Scientifica). The female abdomen was subsequently returned to its initial position to minimize the extent it blocked the male’s field of view.

#### Visually induced courtship

Virgin male flies were collected 2-8h following eclosion and single-housed for 42-48h to increase their motivation to court^11,16,19,22^. All courtship assays were performed at Zeitgeber 0-3h. After plate tethering, male flies were transferred to a humidified warm chamber to recover from anesthesia for 1-3h. Flies were subsequently placed on the center of the ball and left to acclimate for at least 30min in the dark. Each trial was initiated by the presentation of a stationary visual target for 60 sec to examine the animal’s baseline locomotion, after which the visual target began to oscillate. The visual target oscillated in a 75° arc about the animal with a constant angular velocity of ∼75°/s, but the angular size of the dot was continuously altered to mimic the dynamics of a natural female during courtship. The angular size was altered by changing the distance between the male and the target in the ViRMEn world. The distance between the male and the target was taken from the inter-fly-distance (IFD) in a courting pair over the course of two minutes of courtship, and at each frame the angular position of the target was scaled by this IFD to give rise to a more dynamic female path. Angular sizes ranged between −8-50°, with the average size being 22.5°. Each stimulus frame was thus unique for 2 minutes of time, and subsequently repeated until the end of the trial. Experiments ranged from 10-30 minutes in duration, but, for consistency, only the first l0 minutes from each animal was included for further analysis. Across genotypes, - 0 of male flies spontaneously initiated courtship toward the visual target.

#### Optomotor assays

Male flies flies were isolated l2-8h after eclosion and reared with other male flies at low density (5-10 animals) for 2-3 days until adulthood. After plate tethering, male flies were transferred to a humidified warm chamber to recover from anesthesia for 1-3h. Flies were subsequently placed on the center of the ball and left to acclimate for at least 30min in the dark. Wide-field motion stimuli were generated in ViRMEn and consisted of a square-wave grating on a light background with a wavelength of 5° and a rotational velocity of ll5°/s. To allow us to compare neuronal activity during optomotor tracking to baseline periods, we interleaved presentations of a static grating with presentations of a moving grating during each trial, with each epoch lasting for 200s. To approximate the turning responses of courting flies in response to an oscillating target stimulus, the moving grating switched its rotational direction (i.e. clock-wise to counter clock-wise) every 800ms.

### Analysis of behavioral assays

#### Heat maps of turning

Turning was computed on a frame-by-frame basis as the circular distance between the animal’s current heading and the animal’s heading in the next frame. Heat maps were constructed by computing the phase length (in frames) of the stimulus and multiplying it by 3 (3PL). All frames were fit into a matrix of size Nx-3PL. A very small number of remnant frames (typically < 0.5% of frames) at the end of the trial, caused by the total frames not being divisible by 3PLN, were discarded from heat maps but included in all other analysis.

#### Tracking Index

To estimate how well animals were actively tracking the stimulus, we computed the squared Pearson correlation coefficient between the animal’s change in heading and the angular position of the stimulus. The correlation coefficient was calculated using a centered window of 150-300 frames (∼3-5s), except for Fig. 1g-i where correlations were computed in 10 second bins to highlight transitions between courtship and pausing. To compute a tracking index for the optomotor response, we used the correlation between the animal’s change in heading and the change in angular position of the grating stimulus, rather than its absolute angular position.

#### Classification of behavioral epochs

Courtship was classified as any period where the tracking index exceeded 0. (*r*^*2*^>*0*.*4*). To separate periods of running versus periods of courtship (e.g. Fig. 3e) when analyzing responses to visual stimuli we averaged the male’s velocity for three stimulus cycles and selected all periods where the average tracking index was greater than 0.4 versus periods where it was less than 0.4 but the male’s average linear velocity was at least 5mm/s. The threshold value for the tracking index was selected so as to be well above fluctuations in the tracking index during random running. The threshold for velocity was selected to be well above the noise of a fly standing still caused by small vibrations in the floating ball.

#### Response to motion direction

To estimate animal’s behavioral response to motion direction (Extended Data Fig. 8b-e), the stimulus was presented at a slower velocity (80°/s). We selected periods where the stimulus moved progressively on the ipsilateral eye versus periods where the stimulus moved regressively and averaged the turning velocity for each frame across animals.

### Two photon functional imaging

Functional imaging experiments were performed with an Ultima two-photon laser scanning microscope (Bruker Nanosystems) with a Chameleon Ultra II Ti:Sapphire laser. All samples were excited at a wavelength of 920nm, and emitted fluorescence was detected with a GaAsP photodiode detector (Hamamatsu). All images were acquired with a 0X Olympus water-immersion objective with 0.8 NA. To reduce high-frequency noise caused by emitted light from the projector during imaging, we placed a piece of blue-light filter-paper (Rosco) in front of the projector lens, and 3D printed a custom light-shield that fit over the objective to prevent light from entering the brain from above.

After cuticular removal as described above, flies were carefully lowered onto the ball using a micromanipulator (Scientifica) and the shrouded objective lowered over the brain. We subsequently identified the brain region of interest and centered a small ROI over it, yielding an imaging rate of at least 8-l2Hz (except for during P1 activation, where we used larger ROIs yielding imaging rates of -l0Hz). Power was kept low and we ensured that no pixels were saturated. On rare occasions flies were discarded because the glue holding the proboscis broke loose from the mouthparts, causing severe motion artifacts in the *z*-direction.

#### LC10a imaging

An ROI was selected to cover the entire AOTu on a single hemisphere at the depth with the broadest axon terminal distribution, roughly the center of the glomerulus in the superior-inferior axis. Flies were subsequently left to acclimate to the ball for at least 30 minutes before imaging commenced. The hemisphere targeted for imaging was selected pseudorandomly for each fly to ensure that there were no significant biases in expression between the hemispheres. Trials were structured as described under *Visually Induced Courtship*. When temporal specificity was critical (e.g. in comparing LC10a responses to turning responses or characterizing receptive field properties), we used the faster jGCaMP f sensor instead of the brighter GCaMP6s sensor. We observed no differences in the Lgain between experiments using the two sensors (Extended Data Fig. 6). During imaging of LC10a axon terminals in animals expressing GCaMP in all Fru^+^ neurons (Fig. 5), we selected a narrow ROI encompassing the AOTu, avoiding the adjacent LPC due to possible aberrant excitation of P1 neurons expressing CsChrimson as described below. For experiments with two symmetrically opposing targets (Fig 3g-h), animals were allowed to initiate courtship as described under *Visually Induced Courtship*. However, 30 seconds after the first target was introduced, we introduced a second visual target of identical size, whose angular position was equal and opposite to the first. Both eyes thus received identical visual input.

#### P1 imaging

An ROI was selected to cover the lateral junction in the medio-lateral and dorsal-ventral axes, below the P1 arch and above the protrusion of the P1 ring^10^. As with LCl0a imaging, flies were left to acclimate after ROI selection, and the hemisphere targeted for imaging was altered between experiments. Trials were structured as described under *Visually Induced Courtship*.

#### LPC imaging

For spontaneous courtship, the same protocol as for *P1 imaging* was applied. For imaging Fru+ neuropil in the lateral protocerebral complex following optogenetic activation of P1 neruons, we selected a large ROI over the LPC that extended over the AOTu as well, and used a low power output of the laser. This minimized the duty cycle and the imaging power over P1 axons expressing CsChrimson, curbing aberrant excitation^51^. Three lines of evidence suggest that this was sufficient to minimize continuous activation of the CsChrimson channels: (i) animals only occasionally spontaneously initiated courtship towards the visual target, (ii) courtship dynamics were rich and variable with a high frequency of pausing, and (iii) GCaMP6s signals in the LPC still had a significant dynamic range and their fluorescence correlated with courtship (Extended Data Fig. 5). After an ROI had been selected, animals were left to acclimate on the ball for at least 1h to further minimize reverberant excitation in the courtship circuitry caused by P1 activation during dissection and sample placement. During the trial, animals were presented with a single dot oscillating along a ∼75°/s along a 75° arc with constant angular size. During transient stimulation experiments, an LED pulse (l00ms-pulse width at 2.5Hz) was delivered via two l.5-mm optic fibers (Edmund Optics) coupled to two high-power red LEDs (660nm, LED Engin) after 200-300s of baseline recording. The microscope light path was shuttered during the 100ms pulse windows but imaged for 350ms in between pulses to allow us to capture the fluorescence during stimulation. Optic fibers were attached to the objective shroud and fitted immediately above the animal’s head.

To test whether optogenetic activation of P1 neurons could alter the gain of LC10a neurons, we selected an ROI that covered the AOTu but excluded the LPC. After a 60-second baseline period the stimulus target began to oscillate dynamically to drive males to spontaneously initiate courtship. Males were allowed to court the target spontaneously for four minutes, after which we continuously activated P1 neurons optogenetically for six additional minutes as the target continued to oscillate. During continuous P1-activation experiments, we first delivered a single 2 second pulse of red light, and subsequently pulsed the red light (0.2 Hz, l00ms pulse width) and imaged in-between pulses of light.

To estimate visual responses to progressive versus regressive motion, we selected an ROI that covered the AOTu but excluded the LPC. After a 60-second baseline period during which the stimulus target did not move, it began oscillating along the 75° arc, but stopped at each end of the arc for 1 second. This allowed us to better separate responses to progressive versus regressive motion, as they did not immediately follow one-another on the ipsilateral side. As animals typically quickly ceased to court this abnormal stimulus following spontaneous initiation, we continuously activated P1 neurons to ensure a consistent courtship drive.

### Imaging analysis

Image stacks were motion corrected using Non-Rigid Motion Correction (NoRMCorre)^52^ and were subsequently manually validated for motion artifacts. For each experimental recording, an ROI was drawn in FIJI (ImageJ, NIH) across the entire population of interest containing neuropil (e.g. for LCl0a imaging, an ROI was drawn around the bundle of axon terminals in the AOTu) and the average fluorescence extracted. Fluorescence was normalized in MATLAB by assuming that the pre-stimulus/pre-courtship epoch represented the baseline fluorescence of the populations of interest. The average fluorescence of first l00 frames (∼10s) of recording were thus used as the baseline (*F*_*0*_), and a ΔF/F_0_ was defined as: *ΔF*_*i*_ */ F*_*0*_ *=* (*F*_*i*_ *-F*_*0*_) */F*_*0*_, where *i* denotes the current frame. To allow us to compare the shape of the responses of LC10a neuron during courtship and during plain running, we normalized the average response to one stimulus cycle of each animal to its maximum value (i.e. Δ*F*_*i*_ *=ΔF*_*i*_ */ F*_*max*_, Fig 3f).

#### Image-behavior correlations

Because imaging data was collected at a lower frame rate than behavioral data, we downsampled the behavioral data using linear interpolation at the imaging time points to allow us to compute correlations between behavior and imaging. Behavioral data was binned, and the imaging frames corresponding to each bin averaged. To correlate the fluorescence of LCl0a neurons with the fly’s turning (Fig. 3i), we convolved the fly’s heading signal by an exponential with dynamics closely resembling that of jGCaMP f (rise T = l05ms, decay T = l 0ms)^53^ to account for the delay introduced by the calcium sensor. Similarly, to correlate the fluorescence of Fru^+^ LPC neurons expressing GCaMP6s to turning, we convolved the heading signal by an exponential mimicking the dynamics of this slower sensor (rise T = l55ms, decay T = 350ms). When calculating the average evoked response of LC10a to the visual stimulus, we averaged the responses of all stimulus cycles during courtship and running (determined as described above) and found the peak LF/F_0_ of this averaged response for each animal. For computing the gain, we divided the peak-trough difference during running by the peak-trough difference during courting. For estimating the fraction of calcium transients evoking turning (Extended Data Fig. 6g), we found peaks in the calcium signal that were at least 6a greater than the baseline noise when no stimulus was present, and asked whether the animal exhibited an ipsilateral turn (at least 3a greater than the baseline turning) within 500-ms on either side of the peak.

To assess the activity of P1 neurons in the different behavioral epochs of a courtship trial (Fig. 2f), we segregated the data into three categories: periods where animals were actively courting (*r*^*2*^>*0*.*4*), periods where the animal had previously courted and would court at least once more during the trial (‘primed’ periods), and before animals initiated courtship. To ensure that we robustly sampled data from each epoch and not transition periods, only epochs that lasted at least 10 seconds were included in analysis.

To compare the relationship between the responses of visual projection neurons in the AOTu and tracking during spontaneous courtship versus during continuous activation of P1 neurons, we generated 2-D density plots of animal’s tracking indices versus the evoked response of Fru+ neurons in the AOTu. For each stimulus cycle, we computed the averaged tracking index and the maximum LF/F_0_ in the AOTu and counted the number of observations in each bin. This was normalized by the total number of observations across animals.

#### Size-dependence of response

To estimate the extent to which the animal’s turning response and the magnitude of the evoked ΔF/F_0_ of LC10a neurons depended on the angular size of the stimulus, we computed the average angular size of the target stimulus for each stimulus cycle, and computed the average behavioral and neural response for each 2° bin of angular sizes.

### Two photon optogenetic stimulation

#### Activation

For targeted activation of LC10a neurons in *SS1*>*UAS-CsChrimson* animals, animals were placed on the ball and a small “stimulation” ROI over the AOTu was defined as described above. We subsequently identified a second “sham” ROI of similar size, and set the frame rate of both ROIs to l0Hz. The “sham” ROI was adjacent to the “stimulation” ROI and within the same hemisphere, but did not include any fluorescent neuropil. After ROI identification, animals were left to acclimate to the ball for at least 1h before the experiment commenced. During the experiment, we increased the power of the laser to intermediate levels, and switched between focusing the laser over the “stimulation” ROI and the “sham” ROI every - sec (0 frames plus a 750ms delay to switch the laser focus) while recording the animal’s motion in FicTrac as described above. No visual stimulus was presented to the animal. Each trial lasted l0-30 minutes, but only the first l0 minutes were included in analysis for consistency.

#### Silencing

For targeted silencing of LC10a neurons in *SS1*>*UAS-GtACR1*, we selected a “sham” ROI and a “stimulation” ROI as detailed above. Note that only one z-slice of the AOTu was targeted, making two-photon silencing less profound than broad optogenetic silencing but exquisitely spatially targeted, allowing us to focus on one hemisphere without affecting the other. After resting, trials were structured as described under *Visually Induced Courtship*. The laser was focused over the “sham” ROI during the first stimulus oscillations to ensure that animals properly initiated courtship, and subsequently moved between the “stimulation” ROI and the “sham” ROI every 60-90 seconds for the duration of the trial to intermittently silence LC10a neurons in one hemisphere. As with activation, the first l0 minutes of each trial were included in analysis.

### Model of turning dynamics

We constructed a network model of the visuo-motor transformation underlying animal’s behavioral responses to small moving targets during courtship. The core of the model consists of 20 LC10a neurons per hemisphere, modeled as leaky integrate-and-fire units with spike-rate adaptation^54^. Membrane voltage was computer as:

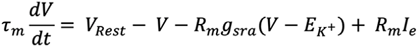

with a membrane time constant (τ_m_)of 100ms, a resting membrane voltage (V_Rest_) of −65mV, a membrane resistance (R_m_) of 10Mn, and a potassium reversal potential (E_K+_) of −70mV. The threshold for spiking is −50mV, after which the membrane voltage is reset to The spike rate adaptation is modeled as a potassium conductance (g_sra_) which increases with each spike in the unit, such that: *g*_*sra*_ *= g*_*sra*_+ *Δg*_*sra*_, where Δ*g*_*sra*_ = 15nS. The potassium conductance is altered such that:

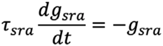

It thus decays to exponentially zero with a time constant τ_sra_ = 200ms. The system of equations was estimated over the entire duration of the trial using Euler’s method with a time-step of 3ms. The input to the model was the angular position of the target stimulus, updated at the same frame-rate as it was presented to animals (−50Hz).

In each hemisphere, each LC10a neuron covers a non-overlapping region of space, ranging from 135° ipsilateral to 15° contralateral (i.e. each neuron covered 15° of the visual field). Each neuron was only sensitive to motion in its designated field-of-view and was generally selective for progressive motion. To compute the input current to each unit of the model, we transformed the estimated receptive fields of LCl0a neurons^23^ into a continuous equation. As the rising phase of the motion-based receptive fields is slower than the falling phase, we fit softmax functions (one rising and one falling) to the two phases independently. We subsequently multiplied these functions to yield a continuous equation for the motion based receptive field. search. Note that the product of the softmax functions is bounded between zero and one, and thus only provided the relative structure of the receptive fields. This relative input strength was scaled by a factor *S*_*f*_ to yield a realistic input current such that:

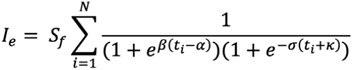

Values κ = 0.8 9, σ = 5.52, α = −0.l86, and β = l5.588 provided the best fit to the estimated receptive field according to a grid search, and we set *S*_*f*_ to 1.5nA. As indicated, the total input current was the sum of the receptive fields up to the current time point, multiplied by the scaling factor. Input was thus primarily driven by visual motion that occurred in the past 500ms, and visual motion that occurred further than ∼2 seconds ago yielded essentially no input current in accord with the estimated receptive fields. Neurons only received input from the time points where the stimulus was present in the neuron’s spatial receptive field.

#### Modelling free behavior

To model turning responses of freely behaving animals in response to female motion, we selected courtship assays during which the male courted the female for at least 90% of the time before copulation. This ensured that our model was primarily compared to the male’s behavior during active courtship. To generate input, we estimated the female position on the male’s retina on each frame as a point as described above and provided this positional information to the model to generate turning. Because the female spent more time in front of the male fly than the target did during open loop, we doubled the visual resolution of the model so that each LC10a model neuron covered only 7.5° of visual space. Because this caused each unit to receive less input current (as the stimulus spent less time within each receptive field), we also increased *S* to 2.5nA to compensate.

To model pursuit in 2-D space (Supplementary Mov. 7), rather than turning responses alone, we initialized the model with the same heading and position as the real male and provided the initial angular position of the female relative to the model. On each frame of the video recording, we updated the angular position of real female fly, and allowed the model to reorient proportionally to the firing rate in the left and right LC10a populations (with a net 200 spikes equaling a 1 radian turn). To move the model in 2-D and allow it to keep up pace with the female, we displaced the model by the same magnitude as the female on each frame, oriented in the direction of the model’s heading. The only exception to this displacement was when the female exhibited an instantaneous velocity less than 2.5mm/s, at which point the model reoriented but was not displaced in 2-D space. Importantly, from this initialization of the model and onwards, the real male and the model were completely decoupled from each other, and independently pursued the target along similar trajectories.

#### Incorporating P1 and LPC activity

To incorporate the fluorescence of P1 neurons and Fru^+^ neurons within the Lateral Protocerebral Complex, we denoised the ΔF/F_0_ time series^55^ and corrected for any drift in the baseline fluorescence using a sliding percentile filter with a window size of 200 seconds. To modulate the response of LC10a neurons in our continuous model, we simply multiplied the input current *l*_*e*_ by the ΔF/F_0_ recorded in the closest previous imaging frame. In our threshold-based model, we set a threshold limit of 0.l5 ΔF/F_0_ (approx. 3a of the baseline ΔF/F_0_ distribution of P1 neurons). When the fluorescence of P1 neurons exceeded this limit, the model received strong input (equivalent of 0.5 ΔF/F_0_ in the continuous model); when it did not, the model received no input on the given frame.

For Fru^+^ neurons imaged with optogenetic activation of P1 neurons, we linearly interpolated the ΔF/F_0_ for the very small number of frames (<20) during which the imaging shutter was closed to allow for optogenetic activation of P1 neurons. Because the peak ΔF/F_0_ often hovered around 0.6 for LPC imaging, we set *S*_*f*_ to 2.5nA to match the peak amplitude (1.5nA) used for modeling continuous tracking when multiplied together with the ΔF/F_0._

### Analysis of model results

To transform spiking in LC10a model neurons to estimates of turning, we implemented a contralateral inhibition component in which we subtract the number of spikes from LC10a neurons in the left hemisphere from LC10a neurons in the right hemisphere in discrete time bins of 30ms. The model turned in the direction with the highest net number of spikes in the given bin, scaled by the magnitude of the net number of spikes. The model was aimed at capturing the dynamics of turning during courtship and the relative magnitude of turns. The absolute magnitude of turning each spike corresponds to in the courting animal is an unconstrained problem and subject to the magnitude of the scaling factor *S*_*f*_. Since *S*_*f*_ was different for tethered behavior and free behavior, and the amplitude of ΔF/F_0_ differed strongly between the LPC signals and P1 signals used for incorporating imaging to the model, we found different estimates for each context: for LPC incorporation we found that a net l00 spikes/s corresponded roughly to l rad/s; for P1 incorporation and during free behavior we found that a net 200 spikes/s corresponded to roughly l rad/s. These values were used for converting spikes to estimated turning (e.g. Fig. 5c).

Each representative alignment of the model and behavior was replicated across at least 4 animals. For computing the Pearson correlation or the cross-correlation between the model’s predicted turning and the animal’s actual turning, we downsampled the model net spiking data using linear interpolation at the behavioral time points since the model frame rate was several times faster than the behavioral recording rate. The animal’s heading signal was smoothed using a moving average window of 30ms. To estimate the fraction of turns that the model accurately predicted (the model “hit” rate) we detected all cycles during which the animal executed an ipsiversive turn of at least 2 radians and calculated the fraction of these peaks that were accompanied by a model in the same direction. To estimate the fraction of turns that the model took but the animal did not take (the model “false alarm” rate), we similarly detected all ipsiversive model turns of at least 1 radian, and calculated the fraction of these peaks that were accompanied by an animal turn of at least 1 radian in the same direction. The threshold for detecting a turn was lower for the model because it, in difference from behavior, is effectively noiseless.

To estimate importance of the temporal structure of the LCl0a receptive fields, we varied the parameter in the input function of our model to yield faster or slower rise-times and compared the Pearson correlation between the model and animals over a 60 sec period of courtship. This variation in the receptive fields causes a narrowing or broadening of the receptive fields, with smaller rise-times yielding less net excitatory input to the model and vise-versa. To correct for this variation, we normalized the area under the curve of the receptive fields by that of the standard model described above (i.e. κ = 0.8 9).

To compute the turning responses of animals and the model for varying levels of P1 activity, we computed the total turning in the direction of the target stimulus for each stimulus cycle, and binned these turning responses based on the maximum ΔF/F_0_ exhibited by P1 neurons in the same stimulus cycle.

### EM connectivity

#### Identifying LC10 subtypes

We analyzed the skeletons and synapses of traced neurons in the female adult hemi-brain connectome^28^ using the Neuprint toolkit in R^56^. 449 LC10 neurons have been labeled as such in the connectome, but the four subtypes (*a-d*) of LC10 neurons^24,57^ have not been separated. To separate these, we manually inspected each of the LC10 neurons and designated each of them as *LC10a, LC10b, LC10c, LC10d*, or *unknown*. We payed particular attention to the somatic location, the branching structure of dendrites in the lobula, the depth of dendritic branches in the lobula, and the branching structure of axons in the large subunit of the AOTu. We were highly selective in our designation and thus denoted a majority of LC10 neurons as *unknown* when we could not ascertain the cell-type based on morphology. We repeated this process thrice and only designated neurons as belonging to a cell-type when it was independently identified on all three rounds. In total, we designated 34 LC10a neurons, 13 LC10b neurons, 21 LC10c neurons, 30 LC10d neurons.

#### Analysis of connectomics data

To group outputs of LC10a neurons by brain region, we examined each of the post-synaptic partners of all LC10a neurons and determined the 3 brain regions in which these partners innervated most densely based on total pre-synaptic counts. To group inputs, we examined each of the pre-synaptic partners LC10a neurons and determined the 3 brain regions in which these partners received the most input based on total post-synaptic counts. Brain regions were defined according to standard meshes included in the hemibrain dataset. We excluded connections from or to optic neuropils, and brain regions to which LC10a neurons were connected to by less than 50 synapses total across the whole population (for example, only one synapse from one LC10a neuron has a post-synaptic partner projecting to the central complex, and this connection was thus not plotted). We plotted the full morphology only of neurons which were strongly connected to LC10a neurons, assessed by the existence of at least 10 synaptic connections between an LC10a neuron and the connected neuron. These were grouped manually as either projecting to the Lateral Accessory Lobes or the Inferior Bridge.

### *Trans*-tango and immunohistochemistry

The *SS1* split-GAL4 line labeling LC10 also labels a sparse number of Kenyon cells^23^, which is amplified to label many Kenyon cells by the *trans*-tango expression system^27^. To visualize the downstream synaptic partners of LCl0a, we instead used a sparser split-GAL4 line that nearly exclusively labels LCl0a neurons (OL00l9B)^24^. This line was not used for functional imaging experiments due to low expression levels. *OL0019B* > *R; Trans-tango* males were selected 8-12h after eclosion and group housed at medium density (l5-20 flies/vial). To allow for proper expression, males were aged for 30-40 days at 20°C before dissections. Brains were dissected in Schneider’s medium (Sigma Aldrich S0146) for 30 min and then immediately transferred to cold 1% paraformaldehyde (Electron Microscopy Sciences) and fixed for l6-20h at °C. Samples were then washed in PAT3 buffer (0.5 BSA, 0.5 Triton X-l00, lX PBS, pH.) three times. The last two washes were made by incubating samples for 1h on a nutator at room temperature. A 3% goat serum block was added, and brains were incubated on a nutator for 90 min at room temperature before block was removed and 1mL of fresh 3% goat serum was added back, along with primary antibodies. Primary antibodies used were 1:50 mouse anti-Brp (nc-82, Developmental Studies Hybridoma bank), l:l000 rabbit anti-GFP (Alll22, Invitrogen; against *OL0019B* > *UAS-myrGFP*), and 1:100 rat anti-HA (11867423001, Roche; against QUAS-mtdTomato). Brains were incubated with primary antibodies on a nutator for 3h at room temperature, and then moved to a nutator at 4°C for 12-16h. Samples were subsequently washed thrice with PAT3 buffer as detailed above. The final wash was removed and 1mL of fresh 3% goat serum added along with secondary antibodies. Secondary antibodies used were 1:500 AF555 goat anti-Rat (A21434, Invitrogen), 1:500 AF633 goat anti-mouse (A21052, Invitrogen), and AF488 goat anti-rabbit (A11034, Invitrogen). Samples were incubated with secondary antibodies for 4 days at 4°C on a nutator, and subsequently mounted in Vectashield (Vector Laboratories) in 5/8^th^ inch hole reinforcements placed on glass slides. Images were captured on an inverted Zeiss LSM 780 confocal microscope using a 25x objective.

### Statistics and reproducibility

P1ease see Supplementary Table l for details on statistics. All statistical analyses were performed in MATLAB R20l9a or GraphPad Prism 8. Data sets were tested for normality using Shapiro-Wilk, and appropriate statistical tests applied as described in Table 1 (e.g. *t*-test for normally distributed data, Fischer’s exact test for categorical observations, Mann Whitney *U*-test for non-parametric data, Friedman test with Dunn *post hoc* test for non-parametric data with repeated measurements). All statistical tests used were two-tailed. Our model was noise-less and thus fully deterministic. Shaded regions surrounding line-plots indicate ± s.e.m. unless otherwise stated. Experimenters were not blind to the conditions of the experiments during data collection and analysis. All attempts at replication were successful.

---

**Extended Data Figure 1.**
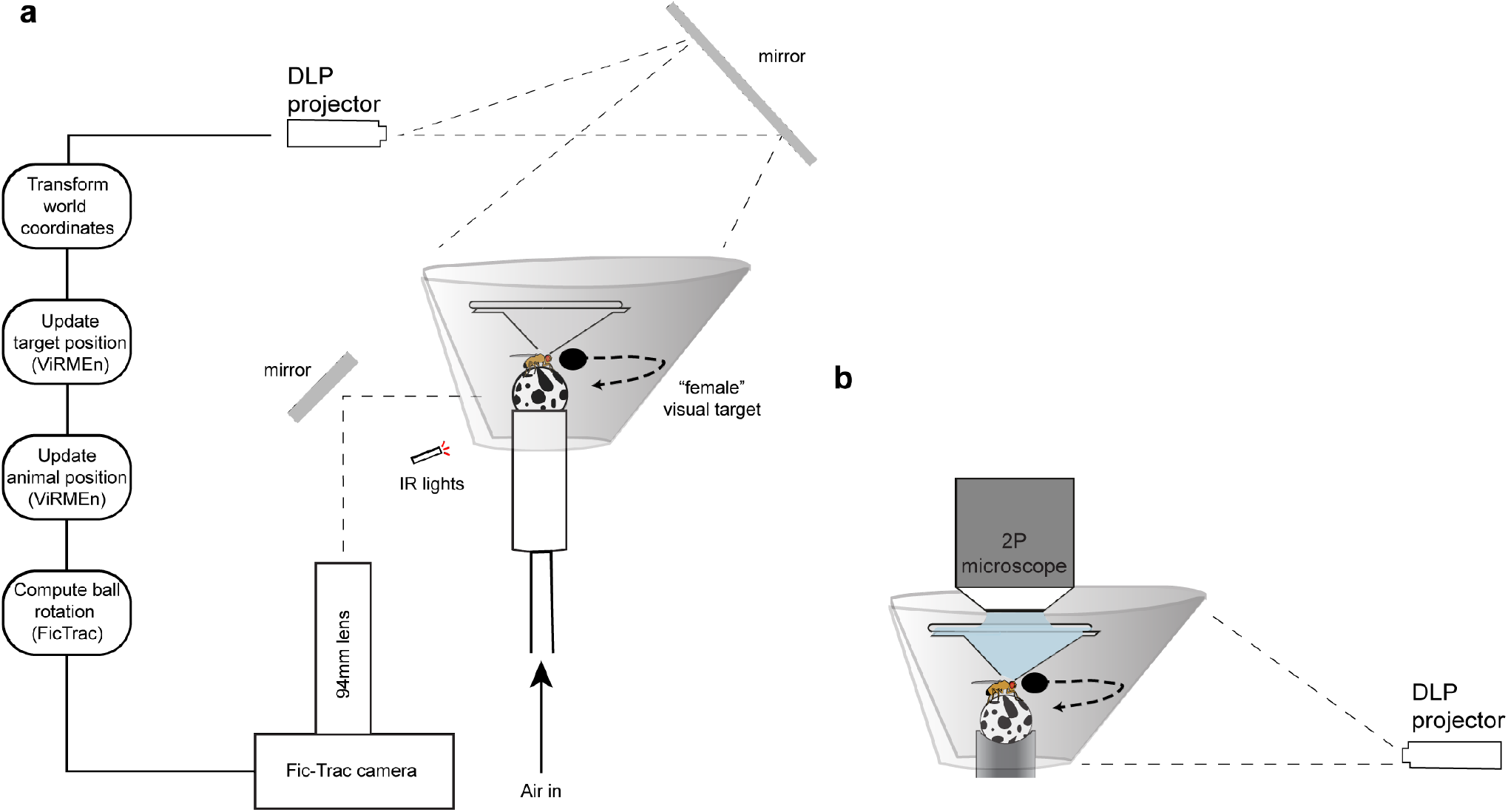
A virtual reality preparation for tethered courtship. **a**, Schematic of virtual reality preparation. Tethered male flies are placed on an air-cushioned foam ball, whose rotational velocity along all three axes is read out by a single camera via the FicTrac software. Based on these rotations, the male position in the virtual world is updated, as is the position of the target stimulus on the screen. Changes in the 2-D world are mapped to a conical screen, and projected by way of a mirror from above. Hardware design from Jazz Weisman and Gaby Maimon. **b**, Schematic of the stimulus presentation during two-photon imaging. Due to sterics of the objective, the stimulus is rear-project onto the screen instead of from above as in (**a**).

**Extended Data Figure 2.**
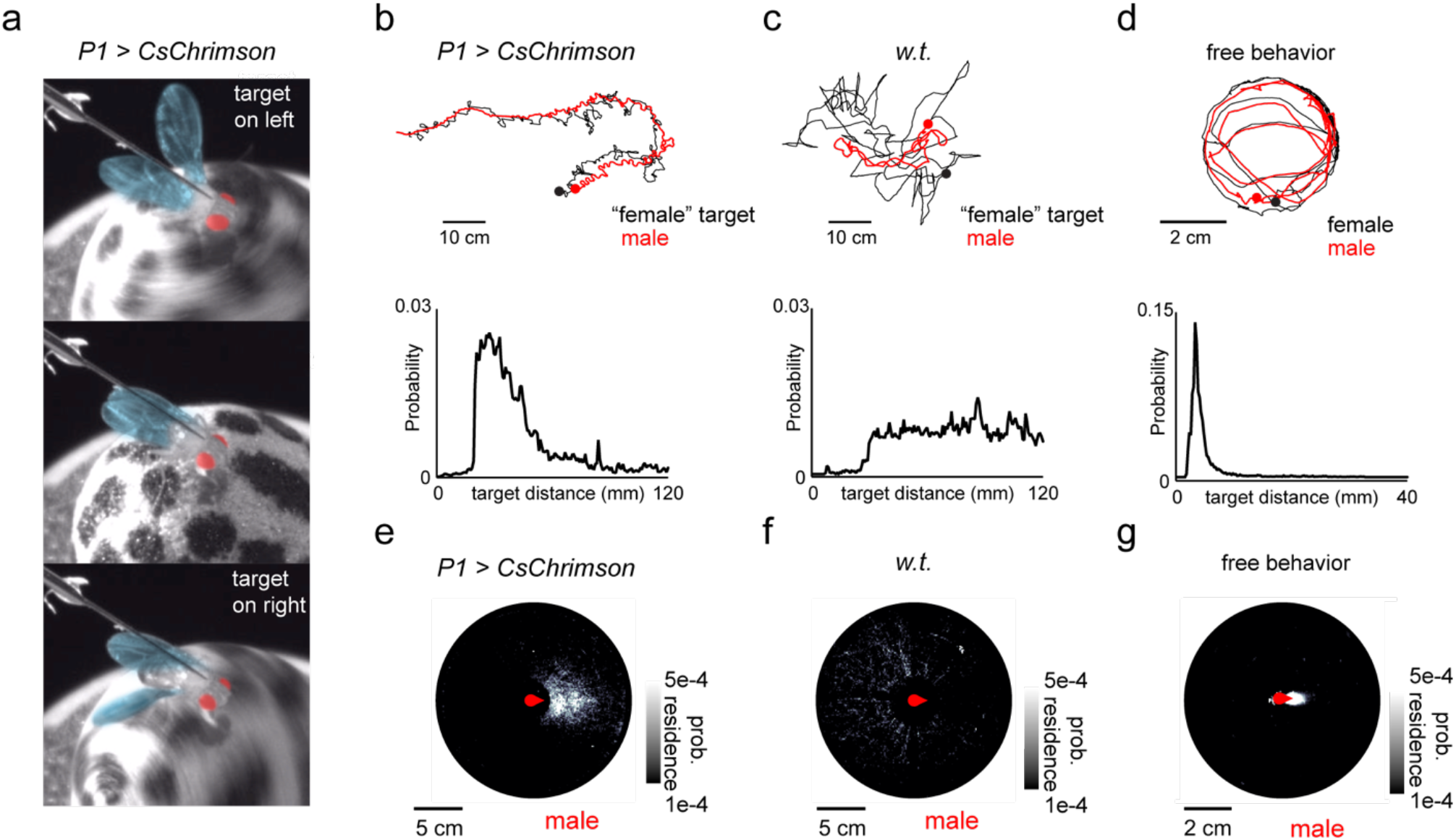
Tethered courtship in an open 2-D virtual world. **a**, Pseudocolor images of a courting male fly during activation of P1 neurons when the visual target is on his left (top), on his right (bottom), or in front of him (middle) showing ipsilateral wing extensions characteristic of courtship song. **b**, Top: position of the male and virtual female in the 2-D world during P1 activation over the course of 200 seconds. Bottom: histogram of the distance between the male and female target in the virtual world during courtship. Note that male is bounded from bringing the stimulus closer than ∼10mm from his position in the virtual world. **c**, Same as (**b**) but for a wild-type male. **d**, Top: Representative example of the 2D positions of the male and female in a freely courting pair of animals. Bottom: histogram of the distance between the male and female fly. **e**, Density plot of the relative position of virtual females with respect to the courting male during P1 activation. **f**, Same as (**e**) but for a wild-type male. **g**, Density plot of the location of the female relative to the male in freely courting pairs of animals. Details of statistical analyses and sample sizes are given in Supplementary Table 1.

**Extended Data Figure 3.**
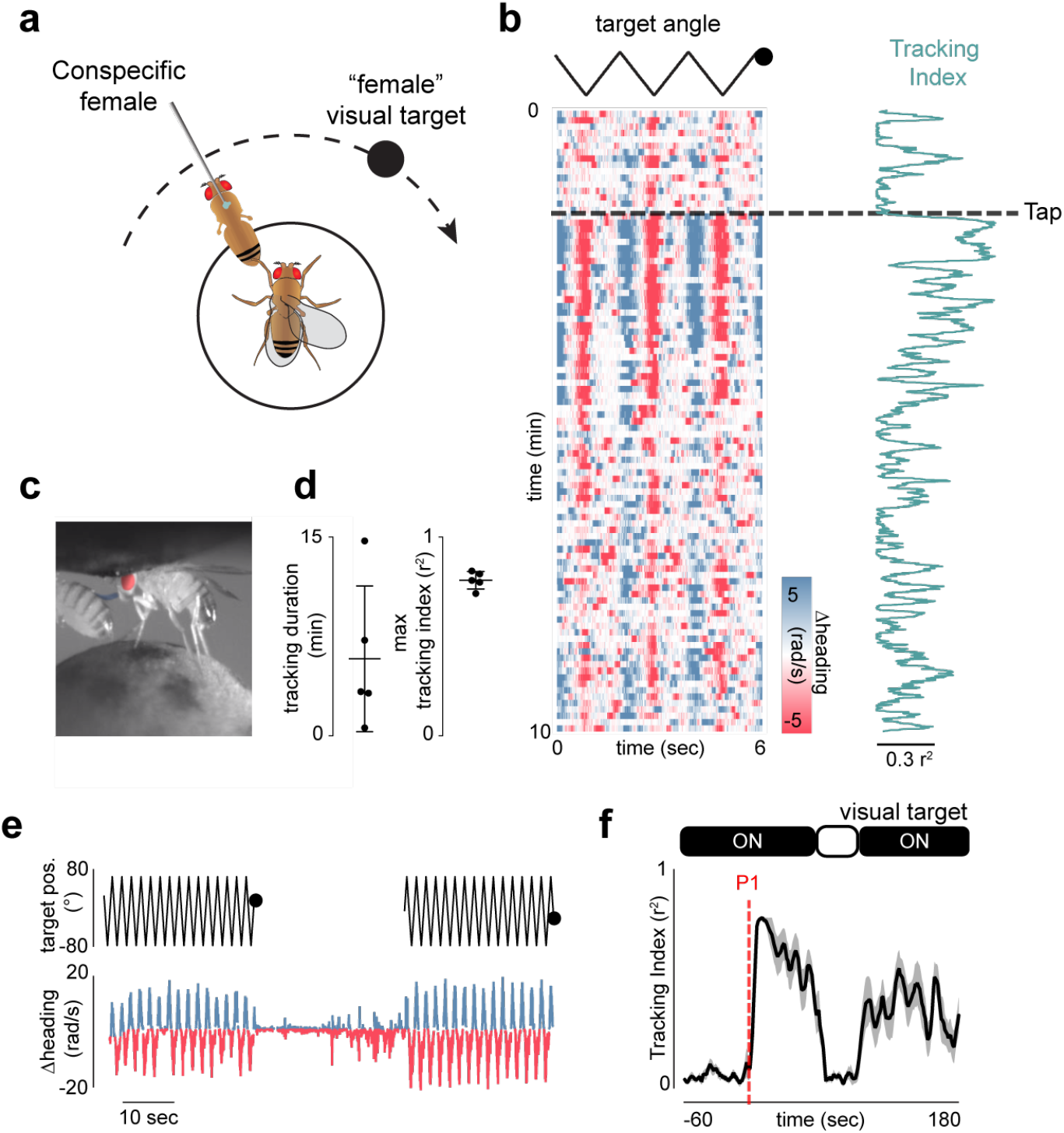
Acute and enduring regulation of courtship arousal. **a**, Schematic of the preparation allowing male to sample gustatory pheromones to trigger courtship. The male fly is provided with the abdomen of a virgin female to taste while the visual target oscillates on the screen in front of him. **b**, Example of a male’s turning during a courtship trial, before and after the male tapped the female abdomen with his foreleg (black line indicates tap). Each row consists of three stimulus cycles. **c**, Pseudocolor image of a male fly sampling the pheromones on a female abdomen. **d**, Maximal tracking index (right) and duration between the first and last detected bout of courtship (left) during pheromone induced courtship trials. **e**, Male orienting during before, during, and after the visual target was transiently removed from the screen (30 s). Courtship arousal was induced by a 3-second optogenetic activation of P1 neurons expressing CsChrimson 60 seconds before stimulus removal. **f**, Average tracking index of males during stimulus-removal trials. P1 neurons were transiently activated 1 min after the visual target began to oscillate, and temporarily removed from the screen 1 min after P1 activation. Details of statistical analyses and sample sizes are given in Supplementary Table 1.

**Extended Data Figure 4.**
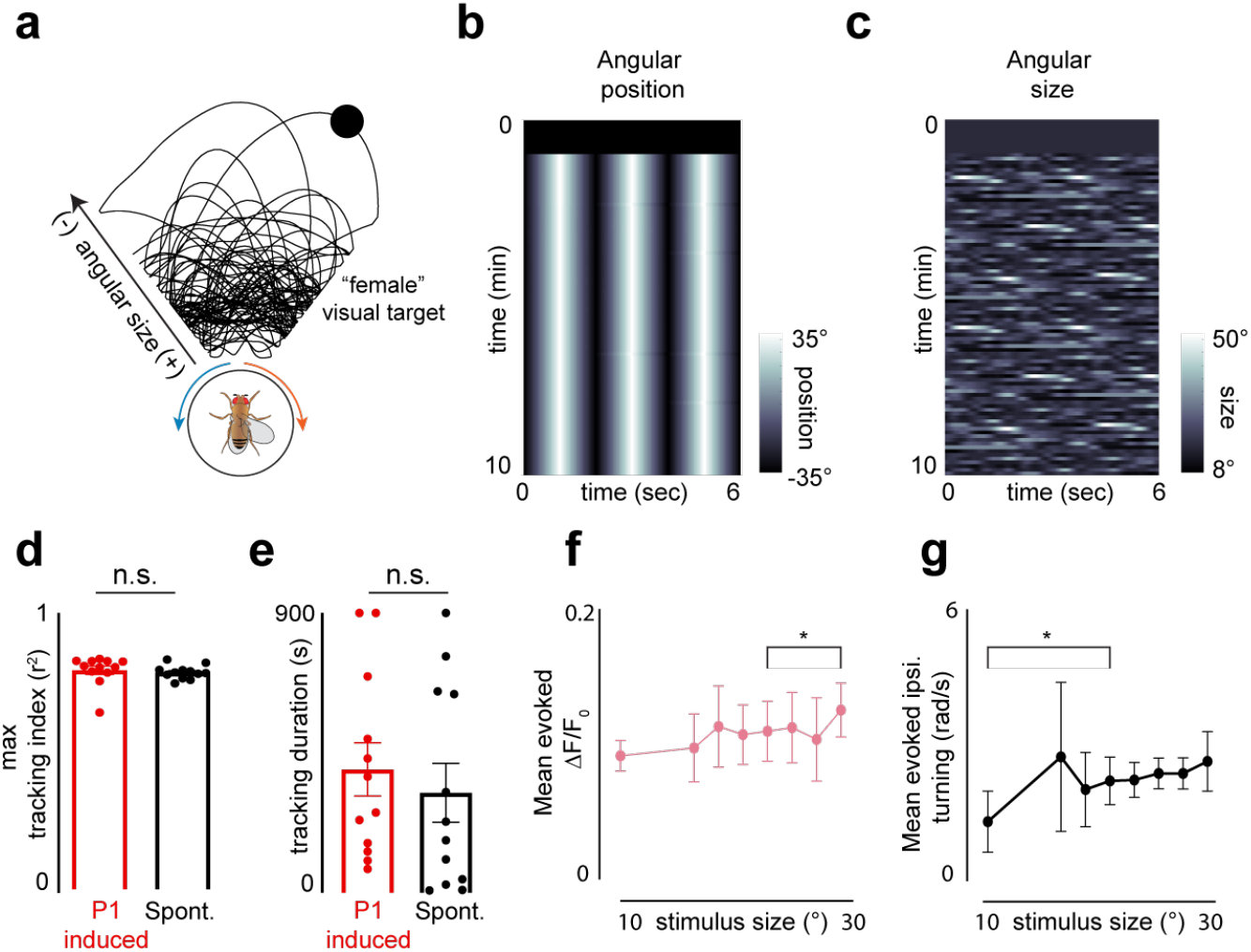
Evoking spontaneous courtship pursuit. **a**, 2-D path of the dynamic visual target used for inducing spontaneous courtship. **b**,**c**, Angular position (**b**) and angular size (**c**) of the dynamic visual target subtended on the male retina over the course of a 10 minute trial. **d**,**e**, Maximal tracking index (**d**) and duration between the first and last detected bout of courtship (**e**) for courtship induced by optogenetic activation of P1 neurons versus spontaneous courtship. **f**,**g**, Average evoked LC10a functional response (ΔF/F_0_, **f**) and average evoked ipsiversive turning (**g**) as a function of the average angular size of the visual target on each stimulus cycle. Details of statistical analyses and sample sizes are given in Supplementary Table 1.

**Extended Data Figure 5.**
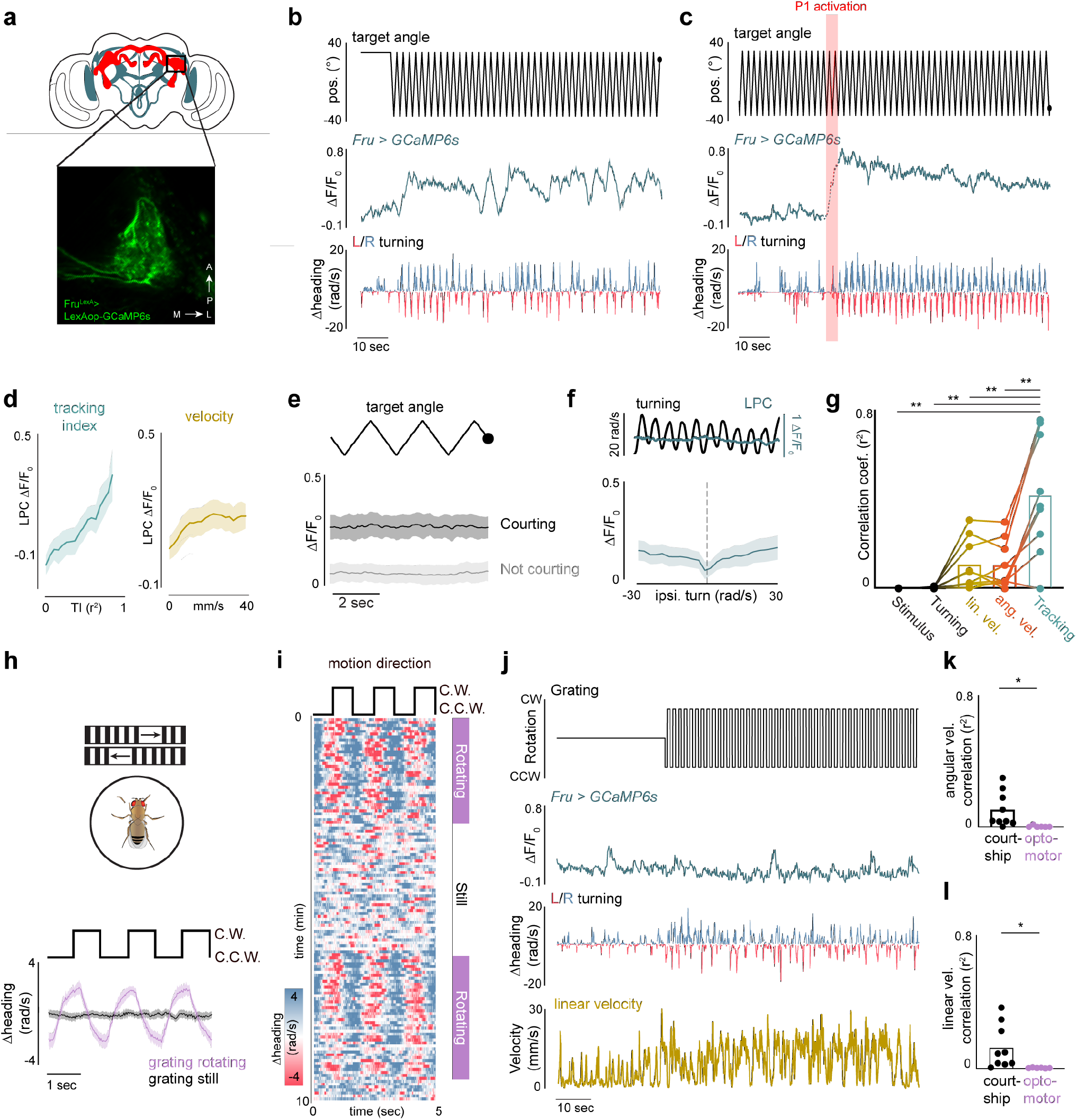
Fru^+^ neurons in the LPC are uncorrelated from the motor implementation of courtship. **a**, Image of Fru^+^ neurons in the LPC expressing GCaMP6s. **b**, Example of the functional responses of Fru+ neuropil in the LPC during a spontaneous courtship trial. **c**, Sample functional response of Fru^+^ neurons in the LPC during an optogenetically induced courtship trial. **d**, Right: functional responses (average ΔF/F_0_) of Fru+ neuropil in the LPC versus behavioral tracking index across all males (r = 0.53). Left: Same as right, but versus the linear velocity of flies (r = 0.09). **e**, Functional responses (average ΔF/F_0_) of Fru^+^ neurons in the LPC during periods of courtship versus periods of random running, aligned to the visual stimulus position. **f**, Top: example of LC10a responses versus male turning. Bottom: Average activity (ΔF/F_0_) of Fru^+^ neurons in the LPC versus the magnitude of ipsiversive turning. **g**, Pearson correlations of the functional responses (ΔF/F_0_) of Fru^+^ neurons in the LPC and the stimulus position, the male’s acute turning responses, the male’s linear velocity, the male’s angular velocity, and the tracking index. **h**, Schematic of preparation for evoking optomotor responses using wide-field motion (top), and the turning responses of animals presented with alternating-direction wide-field motion. **i**, Example of a male’s turning during an optomotor trial. Each row consists of three stimulus cycles. Purple bars indicate when the grating is rotating. **j**, Sample of the functional response (ΔF/F_0_) of Fru^+^ neurons in the LPC during an optomotor trial, before and during the grating oscillates. **k**, Pearson correlations of the functional responses (ΔF/F_0_) of Fru^+^ neurons in the LPC and the animal’s angular velocity during courtship trials or during optomotor trials. **l**, Pearson correlations of the functional responses (ΔF/F_0_) of Fru^+^ neurons in the LPC and the animal’s linear velocity during courtship trials or during optomotor trials. All shaded line plots are mean±s.e.m.; *indicates p < 0.05, **indicates p < 0.01. Details of statistical analyses and group sizes are given in Supplementary Table 1.

**Extended Data Figure 6.**
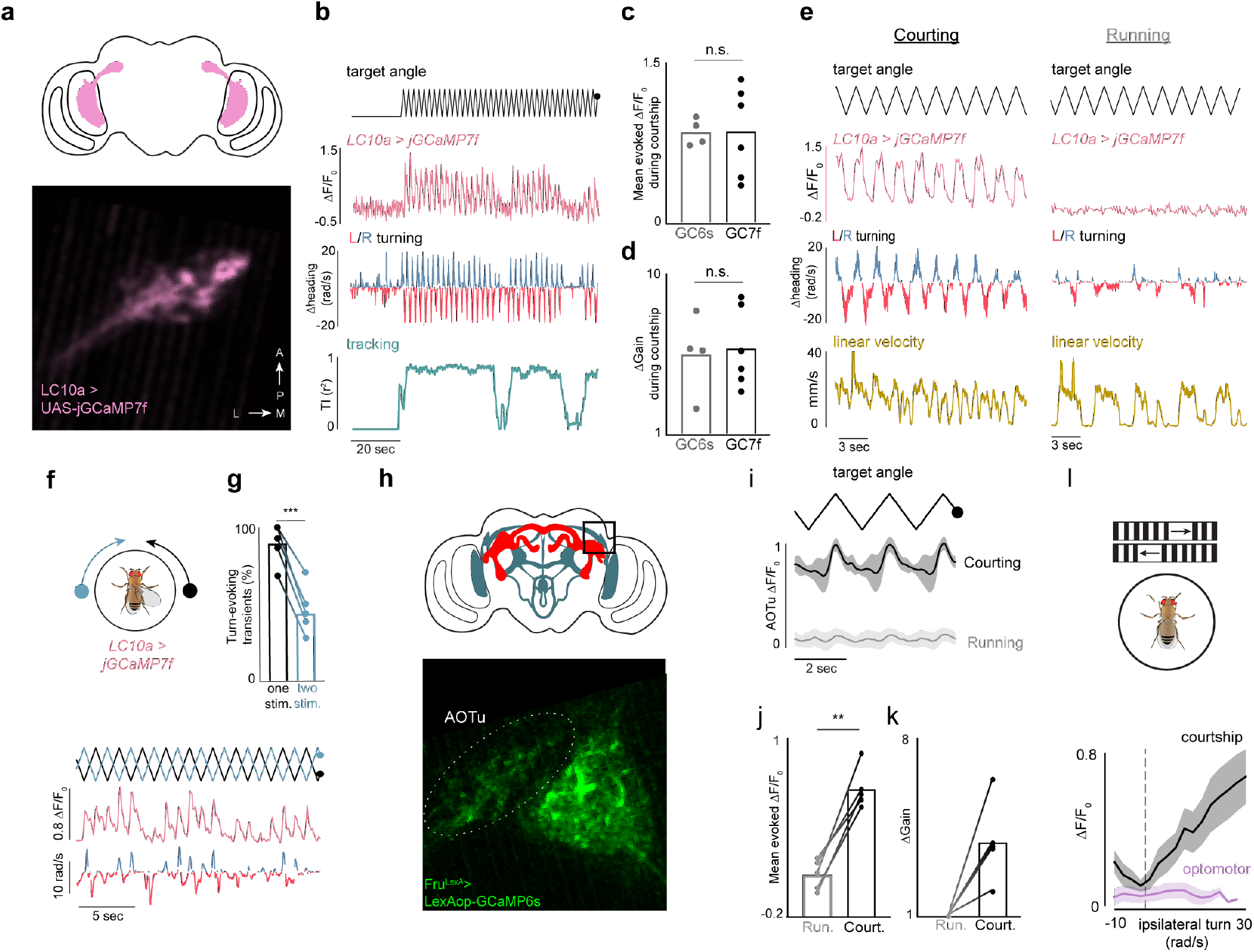
Regulation of LC10a visual responses during courtship. **a**, Image of LC10a axon terminals expressing GCaMP in the anterior optic tubercle. **b**, Example of functional response (ΔF/F_0_) of LC10a neurons expressing jGCaMP7f during a courtship. **c**, Average evoked LC10a responses (ΔF/F_0_) to one stimulus cycle for animals expressing GCaMP6s versus jGCaMP7f. **d**, Average change in LC10a gain (peak-through distance of evoked responses) for animals expressing GCaMP6s versus jGCaMP7f. **e**, Example of LC10a functional responses during courtship versus during a later period of undirected running with similar linear velocity. **f**, Top: schematic of animal being presented with two identical ‘female’ targets, moving in opposition. Bottom: Example of LC10a functional responses versus the position of the two targets (top) and animal turning responses (bottom). Note that LC10a neurons responded even when male failed to turn. **g**, Fraction of calcium transients in LC10a neurons evoking an ipsilateral turning response when animals were presented with one versus two visual targets. **h**, Image of putative LC10a axon terminals in the anterior optic tubercle, expressing GCaMP in all Fru^+^ neurons, and CsChrimson in P1 neurons. **i**, Functional responses (average ΔF/F_0_) of putative LC10a neurons in the AOTu expressing GCaMP driven by Fru-Gal4 during periods of courtship versus periods of undirected running following transient P1 activation, aligned to the visual stimulus position. **j**, Average evoked responses (ΔF/F_0_) of putative LC10a Fru^+^ neurons in the AOTu to one stimulus cycle during courtship versus during periods of random running. **k**, Average change in gain (peak-through distance of evoked responses) of putative LC10a Fru^+^ neurons in the AOTu during courtship relative to during running. **l**, Average activity (ΔF/F_0_) of putative LC10a Fru^+^ neurons in the AOTu versus the magnitude of ipsiversive turning during courtship trials versus during optomotor trials. Details of statistical analyses and sample sizes are given in Supplementary Table 1.

**Extended Data Figure 7.**
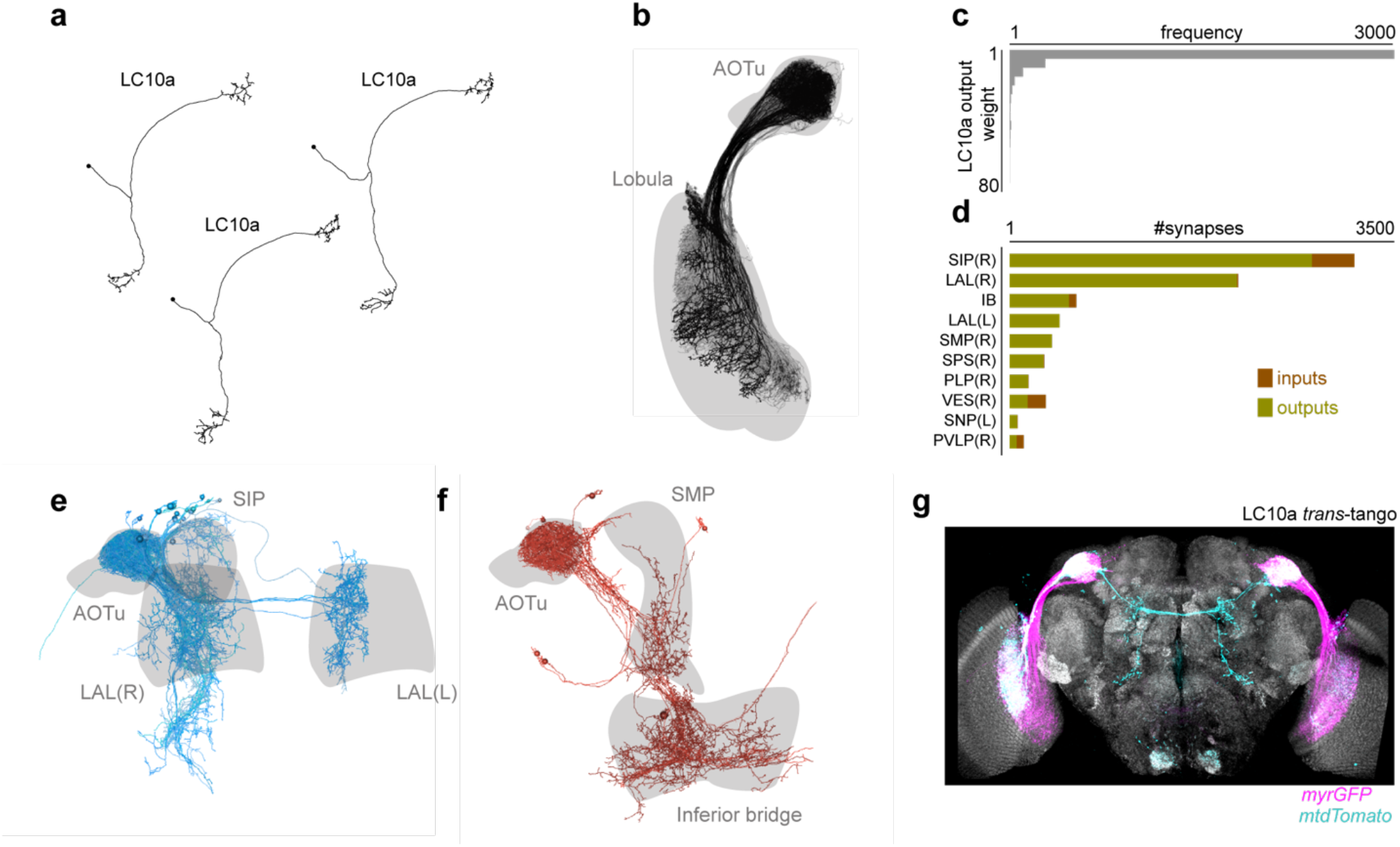
LC10a neurons exhibit sparse and selective connectivity in the central brain. **a**, Examples of identified LC10a neurons in the female hemi-brain connectome^28^. **b**, Morphology of all identified LC10a neurons (n = 34, black) and all other LC10 subtypes (n = 415, grey) in the female hemi-brain connectome. **c**, Histogram of synaptic weights between all LC10a neurons and their post-synaptic partners. **d**, Number of input and output synapses to/from LC10a neurons from the 10 most common brain regions (Superior Intermediate Protocerebrum (SIP), Lateral Accessory Lobe (LAL), Inferior Bridge (IB), Superior Medial Protocerebrum (SMP), Superior Posterior Slope (SPS), Posteriolateral Protocerebrum (PLP), Vest (VES), Superior Neuropils (SNP), Posterior Ventrolateral Protocerebrum (PVLP)), (R/L) indicates the right and left hemisphere, respectively. **e**,**f**, Morphology of all non-optical output neurons from LC10a neurons with at least 10 synaptic connections, grouped based on projections to the LALs (**e**) versus to the IB (**f**). **g**, Representative example of trans-synaptic tracing of LC10a neurons in the male using Trans-Tango^26^. Magenta denotes labeled LC10a neurons, and cyan the labeled post-synaptic partners.

**Extended Data Figure 8.**
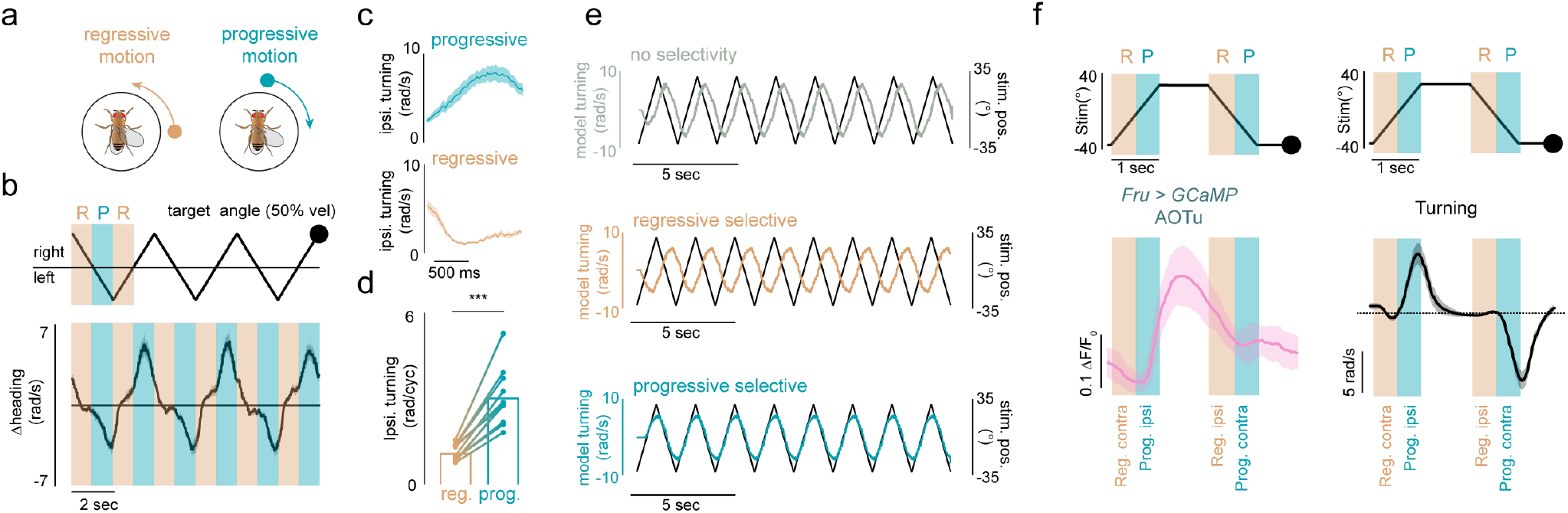
Motion-direction selectivity during courtship pursuit. **a**, Schematic indicating progressive (front-to-back, tan) versus regressive (back-to-front, teal) motion stimuli during tethered courtship. **b**, Average turning responses of a courting male to a slowed-down visual target during continuous optogenetic activation of P1 neurons (teal indicates periods of progressive motion, tan indicates regressive motion). **c**, Average ipsilateral turning of males courting the slowed-down stimulus during periods of progressive versus regressive motion. **d**, Average total ipsilateral turning per period of regressive versus progressive motion of males courting the slowed-down visual stimulus. **e**, Turning responses of LC10a circuit models without motion-direction selectivity, with regressive-motion selectivity, and with progressive-motion selectivity. **f**, Left: average functional response (ΔF/F_0_) of putative LC10a Fru^+^ neurons in the AOTu during optogenetically induced courtship towards a stop-and-go stimulus, separating periods of progressive and regressive motion on the ipsilateral eye by one second. Right: same as left, but for the average animal turning response. Details of statistical analyses and sample sizes are given in Supplementary Table 1.

**Extended Data Figure 9.**
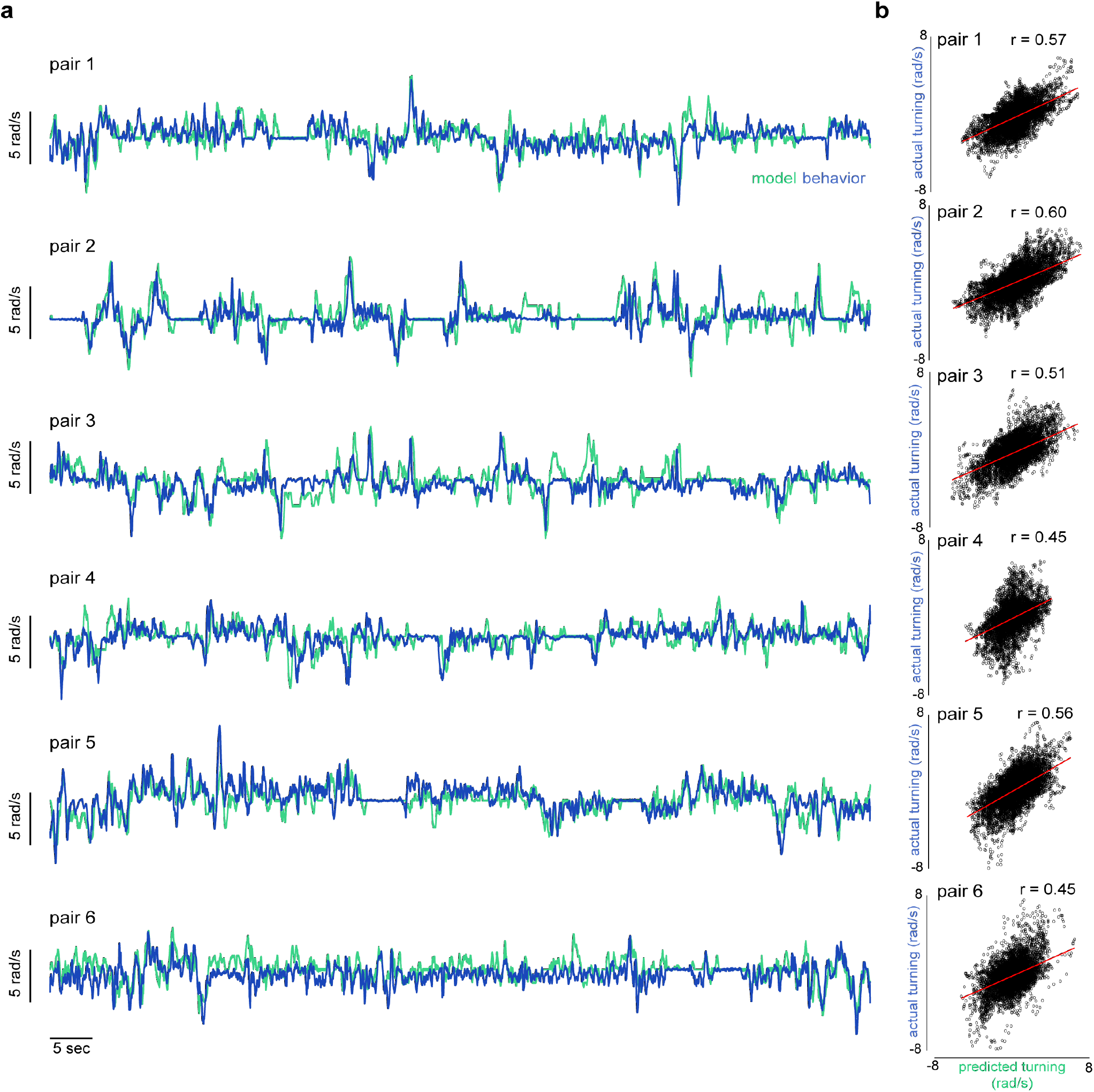
Network model predicts turning dynamics of freely courting males. **a**, Examples of predicted versus actual turning of freely courting males over the first 100 seconds of courtship. **b**, Frame-by-frame predicted versus actual male turning over the course of the full courtship trials for the pairs shown in **a** (5-10 minutes), red line denotes the linear fit. Details of statistical analyses and sample sizes are given in Supplementary Table 1.

**Extended Data Figure 10.**
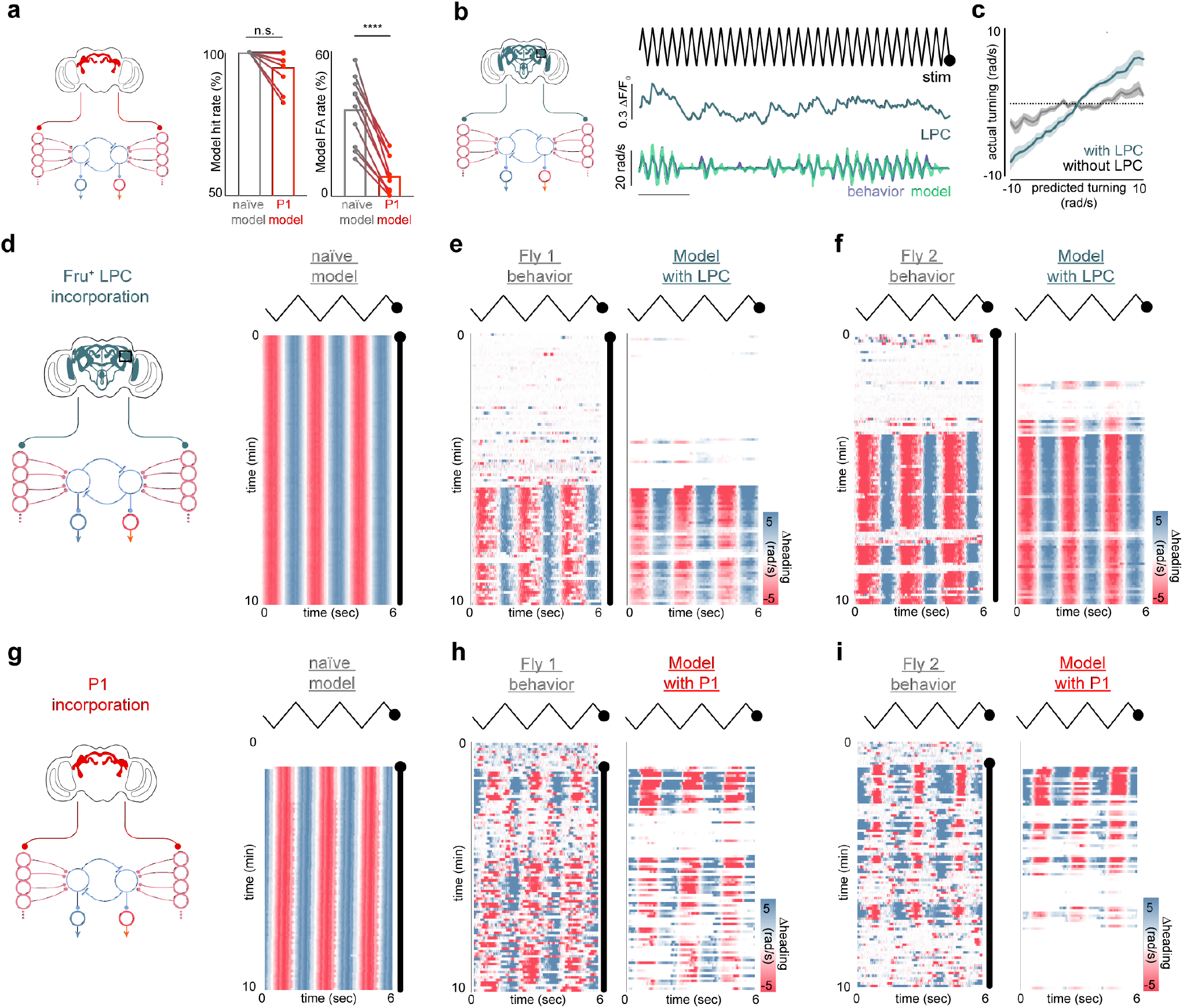
Incorporating P1 or LPC neural activity improves model performance. **a**, Model hit rate (left) and false-alarm rate (right) of the naïve model versus when input current to LC10a neurons is scaled by the functional responses of P1 neurons. **b**, Left: schematic of LPC model – input current is scaled by the instantaneous activity (ΔF/F_0_) of Fru^+^ neurons in the Lateral Protocerebral Complex. Right: example of incorporating the activity of Fru^+^ neurons in the LPC neurons into model for spontaneously courting animals. **c**, Actual versus predicted turning of spontaneously courting animals for models with (blue) and without (grey) incorporated P1 activity. **d**, Example of the predicted turning over a courtship trial by a “naïve-model” (as in Fig. 3a) in which input current to LC10a neurons is consistently high. **e**,**f**, Two examples of actual (left) versus predicted (right) turning responses when the activity of Fru^+^ neurons in the Lateral Protocerebral Complex is incorporated into the model, compare to naïve model in (**d**). **h**,**i**, Two examples of actual (left) versus predicted (right) turning responses when the activity of P1 neurons is incorporated into the model, compare to naïve model in (**g**). Black lines indicate when stimulus is oscillating. Details of statistical analyses and sample sizes are given in Supplementary Table 1.

## Notes

### Competing Interest Statement

The authors have declared no competing interest.

