## Supplementary Movie Legends for "An arousal-gated visual circuit controls pursuit during *Drosophila* courtship"

**Supplemental Movie 1:** Representative example of a male courting a virtual 'female' target in closed loop during continuous optogenetic activation of P1 neurons. Note frequent unilateral wing-extensions, indicating production of courtship song.

**Supplemental Movie 2:** Representative example of the visual stimulus presented to a male courting a virtual 'female' target in closed loop during continuous optogenetic activation of P1 neurons. Female moves autonomously from the male in the virtual world. Note that the male efficiently centers the 'female' target in his field-of-view, and actively brings it closer to him in the virtual world.

**Supplemental Movie 3:** Representative example of a male presented with a simple, repeating visual target before and after P1 neurons are optogenetically activated. Blue line indicates the integrated path of the male. Note the stimulus can be observed as a dark target in the background, and that the male readily exhibits unilateral wing-extensions in the direction ipsilateral to the target after P1 neurons have been activated.

**Supplemental Movie 4:** Example of the translating visual stimulus used to drive males to spontaneously initiate courtship. The target traverses a steady arc but appears to advance and recede.

**Supplemental Movie 5:** Representative example of a male spontaneously initiating courtship towards the translating visual stimulus shown in Supplemental Movie 4. Note that the male does not exhibit any structured turning before the visual stimulus is presented, and the frequent, alternating unilateral wing-extensions, indicating production of directed courtship song.

**Supplemental Movie 6:** Example of the two-target stimulus presented to males, with one target moving slightly slower than the other, causing the phase-relationship between the two to drift over time. 2x playback.

**Supplemental Movie 7:** Three examples of the actual versus predicted paths of how a male pursues a female target in two dimensions. Pink dot is the trajectory of the real female fly; blue dot is the trajectory of the real male fly; green is the trajectory of a simulated fly. The simulated fly is initialized at the same position as the real male fly and matches its linear velocity to that of the real female fly.
