## Supplementary Table 1 for "An arousal-gated visual circuit controls pursuit during *Drosophila* courtship"

### Supplemental Table 1

#### An arousal-gated visual circuit controls pursuit during *Drosophila* courtship

Tom Hindmarsh Sten[1], Rufei Li[1], Adriane Otopalik[1], Vanessa Ruta[1*]

1. Laboratory of Neurophysiology and Behavior, The Rockefeller University, New York, NY, USA.

#### Genotypes

| Male | Male Genotype | Figure |
|---|---|---|
| WT (Canton S) | +; +/+; +/+ | Figs. 1c-d, 3i-k Extended Data Figs. 2c-d, 2f-g, 3b-d, 9a-b |
| Split-P1 > UAS-CsChrimson | w, UAS-CsChrimson.tdTomato; 15A01-AD/+; 71G01-DBD/+ | Figs. 1b-c, 1e, 1g, 1j-k, 3b-h; Extended Data Figs. 2a-b, 2e, 3e-f, 4d-e, 8b-d |
| Split-P1 > UAS-CsChrimson, $Fru^{LexA}$ > LexAop-GCaMP6s | UAS-CsChrimson.tdTomato, LexAop-GCaMP6s; 15A01-AD/Sp; 71G01-DBD/$Fru^{GAL4}$ | Fig. 1f-g, Fig. 5a-e, Extended Data Figs 4d-e, 5a, 5c-l, 6f-k, 8f, 10b-f |
| Fru > UAS-GCaMP6s | w; UAS-GCaMP6s/CyO; Fru-GAL4/TM6B | Extended Data Figs. 5b |
| SS1 > UAS-GCaMP7f | w; VT0012.AD/Sp; VT43656.DBD/UAS-jGCaMP7f | Fig. 1i-k, Figs. 2e-g, Extended Data Figs. 4d-g, 6a-g |
| SS1 > UAS-GCaMP6s | w; VT0012.AD/UAS-GCaMP6s; VT43656.DBD/TM6B | Fig. 1i-k, Fig. 2d Extended Data Fig 4d-e, 6c-d |
| SS1 > UAS-GtACR1 | w; VT0012.AD/+; VT43656.DBD/UAS-GtACR1 | Figs. 2b-c |
| SS1 > UAS-CsChrimson | UAS-CsChrimson.tdTomato; VT0012.AD/+; VT43656.DBD/+ | Fig. 2j |
| OL0019B > *Trans*-Tango | w;R35D04.p65ADZp/UAS-myrGFP, QUAS-mtdTomato; R22D06.DBD/ UAS-hGCG::hlCAM::dNRXN, nsyb-hGCGR::TEVcs::QF, elav-hArr::TEV | Extended Data Fig. 7g |
| Split-P1 > UAS-GCaMP6s | w; 15A01-AD/UAS-GCaMP6s; 71G01-DBD/TM6B | Fig. 1h-k, Fig 2a-i, 5f-g, 5j Extended Data Fig. 4d-e, 10a, 10g-i |

All **females** used (Fig 3j-k, Fig. 1e) were wildtype (+/+; +/+; +/+).

### Statistics and sample sizes

**Fig. 1c**

Analysis: Probability distributions of female positions.

Null hypothesis: The distributions of target angles are equal for wild-type animals and P1>CsChrimson animals.

| Condition | Number of males | Number of trials | Statistical test | Test statistic | P-Value |
|---|---|---|---|---|---|
| W.T. | 9 | 9 | Two-sample Kolmogorov-Smirnov test | 0.24 | <0.000001 |
| P1>CsChrim. | 7 | 7 | | | |

**Fig. 1d**

Analysis: Probability distributions of female positions.

| Condition | Number of male-female pairs | Number of trials |
|---|---|---|
| W.T. | 9 | 9 |

**Fig. 1j**

Analysis: Average turning response to the three cycles of the visual target oscillating in front of the animal for animals where courtship was induced by optogenetic activation of P1 neurons versus for animals where courtship was visually induced.

| Condition | Number of males | Number of trials |
|---|---|---|
| P1 induced | 10 | 10 |
| Spontaneous | 10 | 10 |

**Fig. 1k**

Analysis: Fraction of animals actively engaged in courtship as a function of time since the time of P1 activation or the introduction of the visual target.

Null hypothesis: There is no difference between the distributions of how many animals courted at each moment in time following P1 activation versus following introduction of the visual target.

| Condition | Number of males | Number of trials | Statistical test | Test statistic | P-value |
|---|---|---|---|---|---|
| P1 induced | 10 | 10 | Two-sample Kolmogorov-Smirnov test | 0.1009 | 0.4612 |
| Spontaneous | 10 | 10 | | | |

**Fig. 2d**

Analysis: Average evoked P1 activity and Tracking Index, aligned to the onset of courtship.

| Condition | Number of males | Number of trials |
|---|---|---|
| P1 induced | 7 | 10 |
| Spontaneous | 7 | 10 |

**Fig. 2e, g-i**

Analysis: Correlations between tracking index/visual target position/linear velocity/angular during courtship trials and the $\Delta F/F_0$ of P1 neurons.

Null hypothesis: There is no correlation between the $\Delta F/F_0$ and the parameters during courtship trials animals.

| Figure | Condition | Number of males | Number of trials | Statistical test | Test statistic (mean±st.d) | P-Value |
|---|---|---|---|---|---|---|
| 2e | Tracking Index | 7 | 10 | Pearson's correlation coefficient | 0.562±0.092 | <0.000001 for all animals |
| 2g | Target angle | | | | 0.004±0.006 | > 0.05 for all animals |
| 2h | Linear velocity | | | | 0.231±0.099 | <0.000001 for all animals |
| 2i | Angular velocity | | | | 0.296±0.071 | <0.000001 for all animals |

**Fig. 2e, g-i**

Analysis: Difference in average correlation between the $\Delta F/F_0$ of P1 neurons and the tracking index versus other parameters (visual target position/linear velocity/angular during) courtship trials.

Null hypothesis: The correlation between P1 activity and the tracking index is not different from the correlation between P1 activity and the other parameters.

| Condition | Sample size | Correlation (mean±st.d) | Statistical test | Comparison | Test statistic | P-Value |
|---|---|---|---|---|---|---|
| Tracking Index | 10 trials from 7 animals | 0.562±0.092 | Repeated-measures one-way ANOVA with Geisser-Greenhouse correction, followed by Dunnett's multiple comparison test | N/A | F(9,27) = 3.607 | N/A |
| Target angle | | 0.004±0.006 | | Tracking vs. Target angle | | <0.0001 |
| Linear velocity | | 0.231±0.099 | | Tracking vs. Linear velocity | | <0.0001 |
| Angular velocity | | 0.296±0.071 | | Tracking vs. Angular velocity | | <0.0001 |

**Fig. 2f**

Analysis: P1 activity ($\Delta F/F_0$) before courtship, during active courtship, and during periods where animals were not courting but remained primed to re-initiate.

Null hypothesis: There is no difference in the activity of P1 neurons between the three different behavioral states.

| Condition | Number of trials | Number of samples | P1 activity (mean±st.d) | Statistical test | Comparison | Adjusted p-Value |
|---|---|---|---|---|---|---|
| Pre-courtship | 10 trials from 7 animals | 7213 frames | -0.002±0.09 | One-way ANOVA followed by Tukey's multiple comparison test | Pre vs. Primed | 0.7498 |
| Primed | | 18004 frames | $-2.3\times10^{-6}\pm0.11$ | | Pre vs. Courting | <0.0001 |
| Courting | | 6759 frames | 0.40±0.41 | | Primed vs. Courting | <0.0001 |

**Fig. 3c**

Analysis: Average total turning in the direction ipsilateral to AOTu targeted for silencing during one stimulus cycle.

Null hypothesis: There is no difference between how much animals turn ipsilaterally during sham trials and during silencing trials.

| Condition | Number of males | Number of trials | Value (mean±st.d) | Statistical test | Test statistic | P-Value |
|---|---|---|---|---|---|---|
| Silencing | 3 | 6 | 0.75±0.22 rad/cycle | Student's paired two-tailed t-test | t(5) = 6.526 | 0.0013 |
| Sham silencing | | | 0.29±0.29 rad/cycle | | | |

**Fig. 3e, Left Panel**

Analysis: Average response of LC10a (SS1) neurons expressing jGCaMP7f to the three cycles of the visual target during courtship versus during running without courtship. Conditions are paired for all animals.

| Condition | Number of males | Number of trials |
|---|---|---|
| Courting | 6 | 6 |
| Running | | |

**Fig. 3f**

Analysis: Normalized average response of LC10a (SS1) expressing jGCaMP7f to one stimulus cycle of the visual target during courtship versus during running without courtship. Conditions are paired for all animals.

Null hypothesis: There is no difference in the shape of the two distributions of normalized responses to the visual target during courtship versus during running.

| Condition | Number of males | Number of trials | Statistical test | Test statistic | P-Value |
|---|---|---|---|---|---|
| Courtship | 6 | 6 | Two-sample Kolmogorov-Smirnov test | 0.1557 | 0.2070 |
| Running | | | | | |

**Fig. 3g**
Analysis: Average response of LC10a (SS1) neurons versus turning magnitude in the ipsilateral and contralateral direction.

| Condition | Number of males | Number of trials |
|---|---|---|
| LC10a > jGCaMP7f | 6 | 6 |

**Fig. 3h**
Analysis: Average turning response of animals during two-photon stimulation over the axon terminals of LC10a neurons expressing CsChrimson versus sham stimulation over a random ipsilateral region of similar size.

| Condition | Number of males | Number of trials |
|---|---|---|
| Stimulation + sham | 4 | 4 |

**Fig 4c**
Analysis: Pearson correlation between model and the turning behavior of courting animals in response to a simple oscillating stimulus for various values of κ (rise-time parameter).
Null hypothesis: Varying κ (rise-time parameter) does not affect the correlation between the model and behavior.

| κ value | Number of males | Correlation (r, mean±st.d) | Statistical test | Comparison | Adjusted p-value |
|---|---|---|---|---|---|
| 25% | 5 | 0.77±0.08 | Friedman test | 1v4 | 0.0036 |
| 50% | | 0.83±0.07 | followed by | 2v4 | 0.2126 |
| 75% | | 0.86±0.06 | Dunn's | 3v4 | 0.88 |
| 100% | | 0.89±0.06 | multiple | N/A | |
| 150% | | 0.90±0.01 | comparison | 5v4 | >0.99 |
| 200% | | 0.70±0.08 | test | 6v4 | 0.0036 |

**Fig. 4e**
Analysis: Average turning response of animals to one cycle of the stop-and-go stimulus, compared to the response of the full model, a model without binocularity, and a model not selective for progressive motion.

| Number of males | Number of trials |
|---|---|
| 4 | 4 |

**Fig. 4g**
Analysis: Correlation between animal turning responses and the position of the two targets, as a function of the phase-offset between the two targets.

| Number of males | Number of trials |
|---|---|
| 4 | 4 |

**Fig. 4k**

Analysis: Average cross-correlation between model and the turning responses of freely behaving males during courtship, compared to a shuffled condition where the model responses are scrambled pseudorandomly.

| Number of male-female pairs | Number of courtship trials | Number of simulations |
|---|---|---|
| 6 | 6 | 6 |

**Fig. 5d-e,**

Analysis: Density plot of average tracking index versus evoked $\Delta F/F_0$ of Fru+ projection neurons in the AOTu during spontaneous courtship and during continuous activation of P1 neurons.

| Number of males | Number of trials |
|---|---|
| 6 | **6** |

**Fig. 5h**

Analysis: Moment-by-moment turning of animals versus the turning magnitude predicted by the model at the same moment with and without incorporation of P1 activity.

| Condition | Number of males | Number of trials | Number of simulations | Statistical test | Test statistic (mean±st.d) | P-Value |
|---|---|---|---|---|---|---|
| Tracking Index | 7 | 10 | 10/condition | Pearson's correlation coefficient | r = 0.51±0.11 | <0.000001 for each |
| Velocity | | | | | r = 0.40±0.18 | <0.000001 for each |

**Fig. 5i-j**

Analysis: Average evoked ipsiversive turning response on each stimulus cycle as a function of the $\Delta F/F_0$ of P1 neurons for a continuous model versus a threshold-based model (**l**), or for the continuous model versus the turning responses of animals (**m**),

| Number of males | Number of trials | Number of simulations |
|---|---|---|
| 7 | 10 | 10 / condition |

**Extended Data Fig. 2e-g**
Analysis: Probability residence of the “female” target during the 3D courtship of tethered *P1>CsChrimson* males (**e**), tethered Canton-S wild-type males, and freely behaving Canton-S wild-type males during courtship (**f**).

| Condition | Number of males | Number of trials |
|---|---|---|
| *P1>CsChrimson* | 7 | 7 |
| W.T. (Canton-S) tethered | 9 | 9 |
| W.T. (Canton-S) free behavior | 9 | 9 |

**Extended Data Fig. 3d**
Analysis: Maximum tracking index and duration of courtship for animal’s initiating courtship towards the visual target by tapping the abdomen of a conspecific female.

| Condition | Number of males | Number of trials | Value (mean±st.d) |
|---|---|---|---|
| Tracking duration | 5 | 5 | r = 0.78±0.04 |
| Max tracking index | 5 | 5 | 6.00±5.56 min |

**Extended Data Fig. 3f**
Analysis: the tracking index of animals following a transient (3 second) activation of P1 neurons, with the target stimulus removed for 30 seconds after the first 60 seconds following P1-activation.

| Number of males | Number of trials |
|---|---|
| 9 | 9 |

**Extended Data Fig. 4d-e**

Analysis: Maximum tracking index and courtship duration (time elapsed from first detected courtship bout to the last detected courtship bout)

Null hypothesis: There is no difference in maximum tracking index or courtship duration between optogenetically induced courtship (*split-P1>CsChrimson*) and spontaneously initiated courtship.

| Parameter | Condition | Number of males | Number of trials | Value (mean±st.d) | Statistical test | Test statistic | P-Value |
|---|---|---|---|---|---|---|---|
| Max Track. Ind. | Optogenetic | 12 | | 0.80±0.05 | Student's two-tailed t-test | t(22) = 0.4911 | 0.628 |
| | Spontaneous | 12 | | 0.79±0.02 | | | |
| Courtship duration | Optogenetic | 12 | | 397±296 sec | Student's two-tailed t-test | t(22) = 0.5884 | 0.5623 |
| | Spontaneous | 12 | | 322±326 sec | | | |

**Extended Data Fig. 4f**

Analysis: Average evoked $\Delta F/F_0$ of LC10a (SS1) neurons in response to the dynamic visual target stimulus as a function of its angular size (left)

Null hypothesis: There is no difference in the magnitude of evoked responses when the size of the target stimulus is changed.

| Condition | Number of males | Value (mean±std) | Statistical test | Comparisons | Adjusted p-value |
|---|---|---|---|---|---|
| 28° | | 0.13±0.02 $\Delta F/F_0$ | Repeated-Measures one-way ANOVA with Geiser-Greenhouse correction, followed by Tukey's multiple comparisons test | 28° vs. 26° | 0.5475 |
| 26° | | 0.10±0.03 $\Delta F/F_0$ | | 28° vs. 24° | 0.4224 |
| 24° | | 0.11±0.03 $\Delta F/F_0$ | | 28° vs. 22° | 0.0356 |
| 22° | | 0.11±0.02 $\Delta F/F_0$ | | 28° vs. 20° | 0.0638 |
| 20° | 4 | 0.11±0.02 $\Delta F/F_0$ | | 28° vs. 18° | 0.5849 |
| 18° | | 0.11±0.03 $\Delta F/F_0$ | | 28° vs. 16° | 0.3552 |
| 16° | | 0.10±0.02 $\Delta F/F_0$ | | 28° vs. 10° | 0.0629 |
| 10° | | 0.09±0.01 $\Delta F/F_0$ | | 26° vs. 24° | 0.9943 |
| | | | | 26° vs. 22° | 0.9908 |
| | | | | 26° vs. 20° | 0.9987 |
| | | | | 26° vs. 18° | 0.9479 |
| | | | | 26° vs. 16° | >0.9999 |
| | | | | 26° vs. 10° | 0.9701 |
| | | | | 24° vs. 22° | 0.9986 |
| | | | | 24° vs. 20° | 0.9907 |
| | | | | 24° vs. 18° | >0.9999 |
| | | | | 24° vs. 16° | 0.5896 |
| | | | | 24° vs. 10° | 0.5075 |
| | | | | 22° vs. 20° | 0.9106 |
| | | | | 22° vs. 18° | 0.9795 |
| | | | | 22° vs. 16° | 0.9305 |
| | | | | 22° vs. 10° | 0.4624 |
| | | | | 20° vs. 18° | 0.9362 |
| | | | | 20° vs. 16° | 0.9869 |
| | | | | 20° vs. 10° | 0.5695 |
| | | | | 18° vs. 16° | 0.8873 |
| | | | | 18° vs. 10° | 0.6504 |
| | | | | 16° vs. 10° | 0.9964 |

**Extended Data Fig. 4g**

Analysis: Average evoked turning in the stimulus direction in response to the dynamic visual target stimulus as a function of its angular size (left)

Null hypothesis: There is no difference in the magnitude of evoked turning responses when the size of the target stimulus is changed.

| Condition | Number of males | Value (mean±std) | Statistical test | Comparisons | Adjusted p-value |
|---|---|---|---|---|---|
| 28° | 4 | 3.53±0.88 rad | RM one-way ANOVA with Geiser-Greenhouse correction, followed by Tukey's multiple comparisons test | 28° vs. 26° | 0.8074 |
| 26° | | 3.17±0.46 rad | | 28° vs. 24° | 0.8254 |
| 24° | | 3.18±0.45 rad | | 28° vs. 22° | 0.7546 |
| 22° | | 2.98±0.52 rad | | 28° vs. 20° | 0.8739 |
| 20° | | 2.95±0.71 rad | | 28° vs. 18° | 0.6319 |
| 18° | | 2.80±1.09 rad | | 28° vs. 16° | >0.9999 |
| 16° | | 3.67±2.19 rad | | 28° vs. 10° | 0.2921 |
| 10° | | 1.75±0.90 rad | | 26° vs. 24° | >0.9999 |
| | | | | 26° vs. 22° | 0.8344 |
| | | | | 26° vs. 20° | 0.9744 |
| | | | | 26° vs. 18° | 0.8645 |
| | | | | 26° vs. 16° | 0.9972 |
| | | | | 26° vs. 10° | 0.1972 |
| | | | | 24° vs. 22° | 0.9038 |
| | | | | 24° vs. 20° | 0.9644 |
| | | | | 24° vs. 18° | 0.8468 |
| | | | | 24° vs. 16° | 0.9972 |
| | | | | 24° vs. 10° | 0.1801 |
| | | | | 22° vs. 20° | >0.9999 |
| | | | | 22° vs. 18° | 0.9895 |
| | | | | 22° vs. 16° | 0.9854 |
| | | | | 22° vs. 10° | 0.2471 |
| | | | | 20° vs. 18° | 0.9387 |
| | | | | 20° vs. 16° | 0.9698 |
| | | | | 20° vs. 10° | 0.0172 |
| | | | | 18° vs. 16° | 0.7101 |
| | | | | 18° vs. 10° | 0.2300 |
| | | | | 16° vs. 10° | 0.4691 |

**Fig. 5d**

Analysis: Correlations between tracking index/velocity of animals with the $\Delta F/F_0$ of $Fru^+$ neurons in the Lateral Protocerebral Complex during optogenetically induced courtship.

Null hypothesis: There is no correlation between the $\Delta F/F_0$ and the tracking-index or velocity of animals.

| Condition | Number of males | Number of trials | Statistical test | Test statistic | P-Value |
|---|---|---|---|---|---|
| Tracking Index<br>Velocity | 9 | 9 | Pearson's correlation coefficient | r = 0.53<br>r = 0.09 | <0.000001<br><0.000001 |

**Extended Data Fig. 5e-f**

Analysis: Average response of $Fru^+$ neurons in the LPC to the three cycles of the visual target during courtship versus during running without courtship (**d**), and average activity of $Fru^+$ neurons in the LPC versus turning magnitude in the ipsilateral and contralateral direction (**e**).

| Condition | Number of males | Number of trials |
|---|---|---|
| Courting<br>Running<br>Turning | 9 | 9 |

**Extended Data Fig. 5g**
Analysis: Pearson correlation between the activity of $Fru^+$ neurons in the LPC and instantaneous turning, the stimulus position, the linear velocity of animals, the angular velocity of animals, and the tracking index of animals.
Null hypothesis: The activity of $Fru^+$ neurons in the LPC correlate with other variables as well as it correlates with the tracking index of animals.

| Parameter | Number of males | Correlation (r, mean±st.d) | Statistical test | Test statistic | Comparison | Adjusted p-value |
|---|---|---|---|---|---|---|
| Stimulus | 9 | r=0.001±0.001 | Repeated-Measures one-way ANOVA with Geiser-Greenhouse correction, followed by Dunnett's multiple comparisons test | F(8,32) = 3.11 | Stim vs. Tracking | 0.0052 |
| Turning | | r=0.005±0.004 | | | Turning vs. Tracking | 0.0052 |
| Velocity | | r=0.109±0.03 | | | Linear velocity vs. tracking | 0.0031 |
| Angular velocity | | r=0.105±0.11 | | | Angular velocity vs. tracking | 0.0047 |
| Tracking Index | | r=0.417±0.26 | | | N/A | N/A |

**Extended Data Fig. 5h**
Analysis: Average turning response of animals in response to three cycles of wide-field grating oscillations compared to when animals were presented with a still grating.

| Condition | Number of males | Number of trials |
|---|---|---|
| Grating oscillating | 6 | 6 |
| Grating stationary | | |

**Extended Data Fig. 5j-k**

Analysis: Pearson correlation between the activity of $Fru^+$ neurons in the LPC and the tracking index of animals during courtship versus during optomotor tracking of a wide-field grating (**k**), and Pearson correlation between the activity of $Fru^+$ neurons and the velocity of animals on courtship trials versus during optomotor tracking (**l**).

Null hypothesis: The activity of $Fru^+$ neurons in the LPC correlates with the tracking index and velocity of animals equally during courtship trials and optomotor trials.

| Parameter | Condition | Number of males | Number of trials | Correlation (mean±std) | Statistical test | Test statistic | p-Value |
|---|---|---|---|---|---|---|---|
| Angular velocity | Courtship | 9 | 9 | r=0.105±0.11 | Student's two-tailed t-test | t(13) = 2.296 | 0.0389 |
| | Optomotor | 6 | 6 | r=0.003±0.01 | | | |
| Linear velocity | Courting | 9 | 9 | r=0.109±0.03 | Student's two-tailed t-test | t(13) = 2.303 | 0.0385 |
| | Optomotor | 6 | 6 | r=0.002±0.00 | | | |

**Extended Data Fig. 6c-d**

Analysis: Average evoked $\Delta F/F_0$ (**c**) and Δgain (**d**) of LC10a (SS1) neurons expressing GCaMP7f versus GCaMP6s in response to the visual target stimulus during courtship.

Null hypothesis: The average evoked $\Delta F/F_0$ and Δgain of LC10a (SS1) neurons expressing GCaMP7f and GCaMP6s is not different.

| Parameter | Condition | Number of males | Number of trials | Value (mean±std) | Statistical test | Test statistic | p-Value |
|---|---|---|---|---|---|---|---|
| Evoked $\Delta F/F_0$ | GCaMP7f | 6 | 6 | 0.84±0.12 $\Delta F/F_0$ | Student's two-tailed t-test | t(8) = 0.037 | 0.971 |
| | GCaMP6s | 4 | 4 | 0.85±0.42 $\Delta F/F_0$ | | | |
| ΔGain | GCaMP7f | 6 | 6 | 5.31±2.48 | Student's two-tailed t-test | t(8) = 0.240 | 0.816 |
| | GCaMP6s | 4 | 4 | 4.92±2.52 | | | |

**Extended Data Fig. 6g**

Analysis: Probability that a calcium transient in LC10a neurons evokes an ipsilateral turn for periods when animals are presented with one dot versus two equal and opposite dots.

Null hypothesis: The probability of transients evoking turning is equal for the one-dot and two-dot conditions.

| Condition | Number of males | Number of trials | Value (mean±st.d) | Statistical test | Test statistic | P-Value |
|---|---|---|---|---|---|---|
| One stimulus | 3 | 5 | 89.59±13.08% | Student's paired two-tailed t-test | t(4)=9.823 | 0.0006 |
| Two stimuli | | | 46.12±15.93% | | | |

**Extended Data Fig. 6i**

Analysis: Average response of Fru[+] neurons in the AOTu to the three cycles of the visual target during courtship versus during running without courtship. Conditions are paired for all animals.

| Condition | Number of males | Number of trials |
|---|---|---|
| Courting | 5 | 5 |
| Running | | |

**Extended Data Figs. 6j-k**

Analysis: Average evoked $\Delta F/F_0$ and $\Delta$gain of Fru[+] neurons in the AOTu in response to the visual target stimulus during courtship and during running without courtship.

Null hypothesis: There is no difference in the magnitude of evoked responses or gain during courtship versus during running.

| Parameter | Condition | Number of males | Number of trials | Value (mean±std) | Statistical test | Test statistic | p-Value |
|---|---|---|---|---|---|---|---|
| Evoked $\Delta F/F_0$ | Courting | **5** | **5** | 0.67±0.14 $\Delta F/F_0$ | Student's paired two-tailed t-test | t(4) = 8.15 | 0.0012 |
| | Running | | | 0.09±0.10 $\Delta F/F_0$ | | | |
| $\Delta$Gain | Courting | **5** | **5** | 392±157% | Student's paired two-tailed t-test | t(4) = 4.142 | 0.0032 |
| | Running | | | 100% | | | |

**Extended Data Fig. 6l**
Analysis: Average $\Delta F/F_0$ of $Fru^+$ neurons in the AOTu versus the ipsiversive turning of animals during P1-induced courtship trials versus during optomotor trials.

| Condition | Number of males | Number of trials |
|---|---|---|
| Courtship trials | 5 | 5 |
| Optomotor trials | 6 | 6 |

**Extended Data Fig. 8d-e**
Analysis: Average evoked ipsiversive turning in response to a slowed-down target stimulus during progressive versus regressive periods of motion.
Null hypothesis: There is no difference in the magnitude of evoked turning responses when the motion direction of the target stimulus is progressive versus regressive.

| Condition | Number of males | Number of trials | Value (mean±st.d) | Statistical test | Test statistic | p-Value |
|---|---|---|---|---|---|---|
| Progressive motion | 11 | 11 | 3.04±0.98 rad/cycle | Student's paired two-tailed t-test | t(10)=8.174 | $9.74 \times 10^{-6}$ |
| Regressive motion | | | 1.11±0.29 rad/cycle | | | |

**Extended Data Fig. 8g**
Analysis: Average evoked ipsiversive turning (right) in response to a stop-and-go target stimulus during progressive versus regressive periods of motion, and average evoked response of $Fru^+$ neurons in the AOTu during the same period.

| Number of males | Number of trials |
|---|---|
| 6 | 6 |

**Extended Data Fig. 10a**

Analysis: Fraction of ipsiversive turns that the model correctly predicted when P1 activity was incorporated into the model versus when it was not ("hit-rate", left), and fraction of ipsiversive turns made by the model that were not accompanied by a real turn ("false-alarm" rate, right).

Null hypothesis: There is no difference in the model hit rate or the model false-alarm rate when P1 activity is incorporated versus when it is not.

| Condition | Number of males | Number of trials | Value (mean±st.d) | Statistical test | Comparisons | Adjusted p-value |
|---|---|---|---|---|---|---|
| Hit rate, P1 incorporated | 7 | 10 | 93.73±9.07 % | RM one-way ANOVA with Geiser-Greenhouse correction, followed by Sidak's multiple comparisons test | Hit rate: naïve vs. P1 model | 0.0631 |
| Hit rate, naïve model | | | 100±0 % | | | |
| False-alarm rate, P1 incorporated | | | 8.03±7.59 %XW | | FA rate: naïve vs. P1 model | <0.0001 |
| False-alarm rate, naïve model | | | 35.87±14.26 % | | | |

**Extended Data Fig. 10b-c**

Analysis: Acute animal turning as a function of model turning when the model incorporated the activity of $Fru^+$ neurons in the Lateral Protocerebral Complex versus when it did not (naïve model).

| Condition | Number of males | Number of trials | Number of simulations |
|---|---|---|---|
| With $Fru^+$ LPC activity | 9 | 9 | 9 |
| Without $Fru^+$ LPC activity | | | 9 |
